# Controlled exchange of protein and nucleic acid signals from and between synthetic minimal cells

**DOI:** 10.1101/2022.01.03.474826

**Authors:** Joseph M. Heili, Kaitlin Stokes, Nathaniel J. Gaut, Christopher Deich, Jose Gomez-Garcia, Brock Cash, Matthew R. Pawlak, Aaron E Engelhart, Katarzyna P. Adamala

## Abstract

Synthetic minimal cells are a class of small liposome bioreactors that have some, but not all functions of live cells. Here, we report a critical step towards the development of a bottom-up minimal cell: cellular export of functional protein and RNA products. We used cell penetrating peptide tags to translocate payloads across a synthetic cell vesicle membrane. We demonstrated efficient transport of active enzymes, and transport of nucleic acid payloads by RNA binding proteins. We investigated influence of a concentration gradient alongside other factors on the efficiency of the translocation, and we show a method to increase product accumulation in one location. We demonstrate the use of this technology to engineer molecular communication between different populations of synthetic cells, to exchange protein and nucleic acid signals. The synthetic minimal cell production and export of proteins or nucleic acids allows experimental designs that approach the complexity and relevancy of natural biological systems.

## Introduction

In 1932, the surgeon George W. Crile Jr. published methods for creating model cell-like structures from complex organic substances that were able to mimic certain cellular tasks, often cited as the first synthetic cells ^1^. Today, researchers are taking a two pronged approach to creating synthetic cells: top-down and bottom-up^2,3^. The top-down approach involves trimming down the genome of a living cell to the fewest number of essential genes, while the bottom-up approach assembles synthetic cells from non-living basic components ^3,4^. There are many definitions of synthetic cells, but the common trait is that each class of synthetic cell is a simplified mimic of highly complex processes existing in live cells^5,6^. Despite being a relatively new field, the potential and realized applications for this technology are vast. Synthetic cells are being studied for use as bioreactors in bio catalysis ^7^, optimization of protocell models ^8^, modeling for mechanisms of cellular Darwinian competition ^9^, in medical applications such as gene or drug delivery ^5,10^, and in biosensing ^11,12^. Because synthetic cells offer such an important avenue for cellular biology research, there is a focus on modeling the complex processes of naturally occurring cells in synthetic mimics^6^. Cell-like bioreactors containing only the components of interest allow research on a single cellular process without interference from confounding variables often present in cellular experiments. The increasing ability of synthetic minimal cells to bridge the gap between *in-vitro* and *in-vivo* experiments provides a method to investigate foundational questions of cellular function.

A major hurdle in the expansion of synthetic cell capabilities is the export of a product to the outer environment, or the incorporation of a signal from outside the cell. Export of protein or nucleic acid products synthesized within a liposome offers the ability to create artificial cell signaling molecules or facilitate membrane anchoring of extracellular receptors^13^. Import of similar molecules could unlock a caged function, modulate a gene circuit, or provide a step in a modeled metabolic pathway.

Cell penetrating peptides (CPP) have shown promise as membrane-translocating chaperones. Here, we demonstrate a method to mimic cellular import or export of proteins within a synthetic cell. We sought to determine whether CPP can carry a protein cargo across the membrane of a synthetic minimal cell, and verify the efficacy of the cargo post translocation.

## Results and discussion

Throughout this work, we define a synthetic cell as a liposome bilayer membrane with an *E coli* cell-free protein expression system (TXTL) encapsulated inside. That synthetic cell is capable of synthesizing protein from plasmid DNA templates. In most cases used in this work, transcription is catalyzed by T7 RNA polymerase, which we add to the synthetic cells as purified protein (overexpressed in *E coli*).

To investigate whether proteins synthesized in synthetic cells can translocate across a phospholipid bilayer membrane, we investigated five well characterized cell-penetrating peptide (CPP) tags. We tested tags that were previously demonstrated to translocate through bilayer membranes in vitro. The tested tags were: Transportan 10 (AGYLLGKINLKALAALAKKIL)^14^, RL9 (RRWWRRWRR)^15^, FSKRGY (RRGFSLSLALAKDGWALMLRLGYGRR)^16^, penetratin (RQIKIWFQNRRMKWKK)^17^ and Pept 2 (PLIYLRLLRGQF)^18^. We created a fusion protein containing the tested CPP tag, GGGSGGGS spacer and eGFP, using UTR and terminator optimized for E coli cell-free TXTL^19^.

We prepared synthetic cells containing the cell-free protein expression system with 10nM of each of the tested plasmids. The synthetic cell membranes was labeled with lipid-conjugated red fluorescent dye Rhodamine (**Figure 1a** and **b**). Each sample was incubated at 30°C for 16 hours, and then purified on a size exclusion column to separate the liposomes from the unencapsulated solutes (**Figure 1c-g**). The position of liposomes was indicated by red fluorescence of rhodamine membrane dye. Total fluorescence of all liposome fractions and all free solute (outside of liposome) fractions was measured. In those experiments, approximately 7% of total sample volume was the liposome lumen (see Supplementary information for vesicle lumen volume calculations). Since we expect all tested CPP tags to facilitate concentration gradient driven membrane translocation, we repeated those experiments increasing the outside volume so that approximately 3.5% of total volume was inside synthetic cells. **Figure S1** shows all GFP fluorescence results with all tested CPP tags. As expected, in both sets of experiments, in the absence of CPP tags all synthesized protein remains inside vesicles (so all measurable fluorescence comes from vesicle fraction, with minimal free solute fraction fluorescence). We identified two CPP tags that produced the most translocation of GFP payload across the membrane: penetratin and RL9, with penetrating producing significantly more translocation. We selected penetratin tag for further studies.

**Figure 1.**
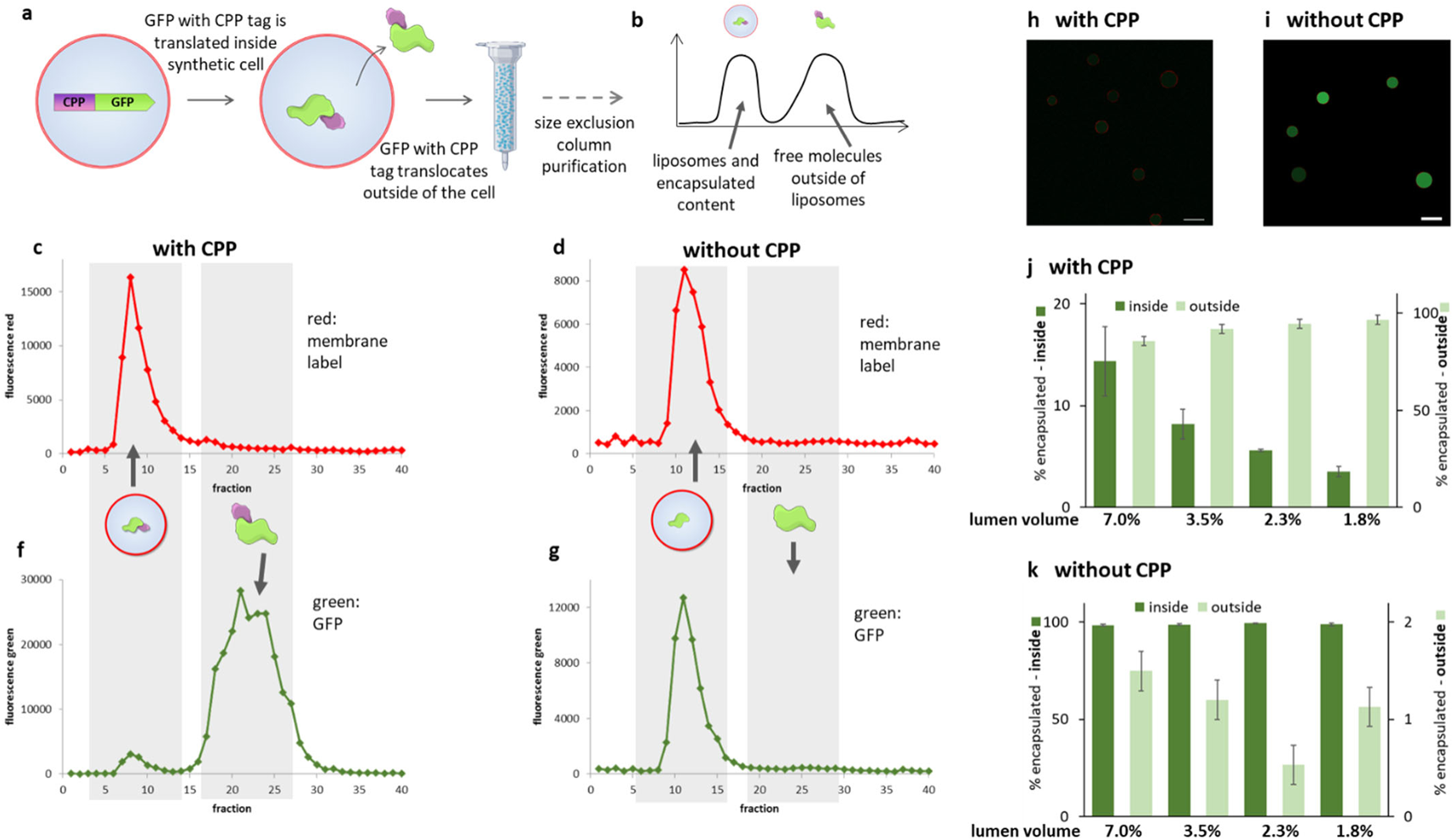
Cell-penetrating peptide labeled GFP crosses synthetic cell vesicle membrane. **a**: General scheme of the experiment. Synthetic cell membrane made from POPC and cholesterol with red Rhodamine dye encapsulating a cell-free translation system and a plasmid for GFP with a CPP label. The CPP-GFP is synthesized and then translocates outside of the cell. **b**: The sample is purified on a size exclusion column, separating liposomes from free solute. **c** and **d**: Example red fluorescent traces during purification of synthetic cell sample, indicating fractions containing the synthetic cell liposomes. **f** and **g**: Example green fluorescent traces during purification of synthetic cell sample, indicating fractions containing GFP. **c** and **f**: samples with CPP tagged GFP. **d** and **g**: samples with untagged GFP. **h**: Fluorescent microscopy image of synthetic cell samples containing CPP tagged GFP. **i**: same as **h** but with untagged GFP. **j** and **k**: Percent of GFP encapsulated inside (left Y axis) and percent of GFP outside the cell (right Y axis) for samples containing different amount of synthetic cells, corresponding to different % of the total sample volume being encapsulated inside the vesicles. See **Table S1** for liposome concentrations corresponding to different lumen volume. **j**: with CPP tagged GFP, **k**: with untagged GFP. Error bars are S.E.M, n=3. The data presented here are for 5nM GFP plasmid, data for experiments with lower plasmid concentration are on **Figure S3**.

To further characterize the performance of penetratin CPP tag, we prepared samples where the synthetic cell lumen was an even smaller fraction of total sample: 2.3% and 1.8%. Comparing GFP fluorescence inside and outside of synthetic cells with decreasing internal volume as a percentage of total sample volume, we note that the fluorescence inside synthetic cells scales proportionally with decreasing internal volume, but never matches the percent internal volume (**Figure 1j**). For example, when approximately 7% of the total sample volume is inside liposomes, 14.4% of total sample GFP fluorescence comes from the vesicle fraction. Similar is true for all other tested lumen to total sample ratios. This might mean either the CPP tag does not cause complete equilibration across the membrane, the vesicle fraction co-purifies with some GFP stuck to the surface of the membrane, or is caught mid-translocation. The GFP fluorescence inside vesicles does decrease as the relative external volume increases, which indicates that the CPP tag facilitates translocation driven by the concentration gradient, as expected.

Negative control of samples without CPPs confirm that CPPs are necessary for GFP translocation – 98% or more of total GFP fluorescence is contained in vesicle fractions for samples with GFP without CPPs, for all lumen to total sample ratios (**Figure 1k**).

Fluorescent microscopy images of samples containing either GFP with (**Figure 1h**) or without CPPs (**Figure 1i**) confirm that the vesicles remain spherical and in absence of CPPs, all GFP fluorescence stays inside synthetic cells.

Next, we investigated whether the CPP tag can facilitate translocation of a bigger, catalytically active protein: T7 RNA polymerase (T7 RNAP). Synthetic cells expressing CPP-tagged T7 RNAP were incubated with DNA template for the fluorescent RNA aptamer broccoli^20^, and all components of the in-vitro transcription reaction added outside of synthetic cells (**Figure 2a**). Another set of samples was prepared, with broccoli template inside synthetic cells. In samples with CPP-tagged T7 RNAP, broccoli fluorescence was observed both when the broccoli template was inside as well as when it was added outside of synthetic cells, indicating that T7 RNA polymerase can catalyze transcription after translocating outside of synthetic cells (**Figure 2b, c** and **d**). In samples with T7 RNAP without a CPP tag, fluorescence was observed only when the broccoli template was inside synthetic cells, and no fluorescence was observed when the broccoli template was added only to the outside of cells (**Figure 2b, e** and **f**). This indicates that T7 RNAP cannot cross the synthetic cell membrane without CPPs, and neither can broccoli template cross the membrane on its own – since in absence of a CPP tag, all T7 RNAP stays inside, and no fluorescence is observed if broccoli template is only outside.

**Figure 2.**
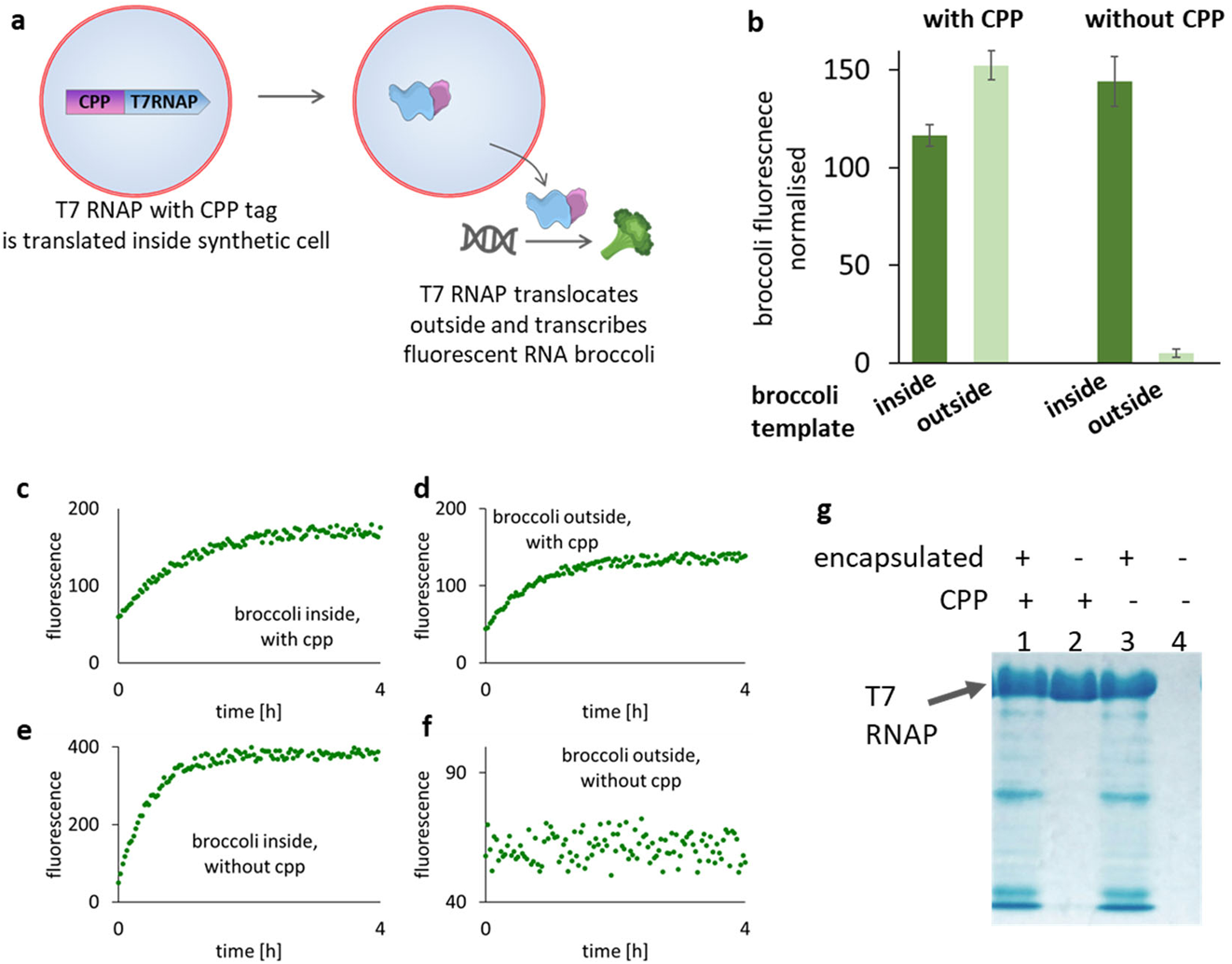
The CPP tag facilitates export of active RNA polymerase. **a**: Schematic of the experiment: synthetic cell with a red labeled membrane expresses T7 RNA polymerase with CPP tag. The T7 RNAP translocates outside of cell, where it catalyzes synthesis of fluorescent RNA broccoli from the DNA template. **b**: Florescence of broccoli RNA aptamer measured inside and outside of synthetic cells, after a 16h reaction followed by purification of samples on a size exclusion column. The “with CPP” and “without CPP” designations refer to the CPP tag on T7RNAP. The broccoli DNA template was either added outside of synthetic cells, or encapsulated inside synthetic cells. Example size exclusion purification traces are on **Figure S4**. **c**-**f**: time courses of broccoli RNA aptamer fluorescence during synthesis. **c** and **e**: broccoli template inside synthetic cells, **d** and **f**: broccoli template outside, **c** and **d**: T7RNAP with CPP tag, **e** and **f**: untagged T7RNAP. **g**: Western blot analysis of T7RNAP from fractions encapsulated inside (+) and outside (-) synthetic cells, with and without CPP tag on the T7RNAP.

Penetratin translocates across the membrane without forming a pore.^15,21^ We hoped that this mode of translocation will ensure that the CPP tag does not induce non-specific membrane leakage in synthetic cells.

To test this, we designed an experiment where synthetic cells express CPP-tagged firefly luciferase. Synthetic cells membranes are labeled with a red dye (rhodamine), and a small molecule membrane-impermeable dye (calcein) was added to the outside of synthetic cells. (**Figure 3a**). After a 16 hour incubation, samples were purified on a size exclusion column, separating the vesicle fraction (identified by red fluorescence of membrane dye) from the free solute fraction (**Figure 3b** and **c**). We lysed vesicles with 0.1% v/v Triton, and as a control used the same amount of Triton in the free solute fraction. We measured the fluorescence of calcein and the activity of luciferase. When calcein dye was added to the outside of synthetic cells, no calcein was detected inside, in the vesicle fraction (**Figure 3f**). This was true in control samples with fluorescein without a CPP tag as well (**Figure 3g**). Luciferase activity measured in samples with CPP-labeled luciferase followed previously observed CPP-labeled GFP distribution: most of the luciferase ended up outside of the synthetic cells (**Figure 3d**). In control samples where luciferase was not CPP-tagged, nearly all luciferase activity was inside synthetic cells, not outside (**Figure 3e**). We reversed the experimental design, adding calcein inside synthetic cells instead of outside. The calcein fluorescence measured after incubation and purification shows that calcein remains inside synthetic cells, both in samples with CPP-tagged luciferase (**Figure 3j**) as well as in samples without CPP-tagged luciferase (**Figure 3k**). The luciferase activity in those experiments also followed the predicted distribution: when luciferase was CPP-tagged, more luciferase ended up outside of synthetic cells (**Figure 3h**), and when luciferase was not CPP-tagged, it remained inside synthetic cells (**Figure 3i**). This demonstrates that presence of a CPP tag translocating protein payload does not make the synthetic cell membranes leaky to small molecules.

**Figure 3.**
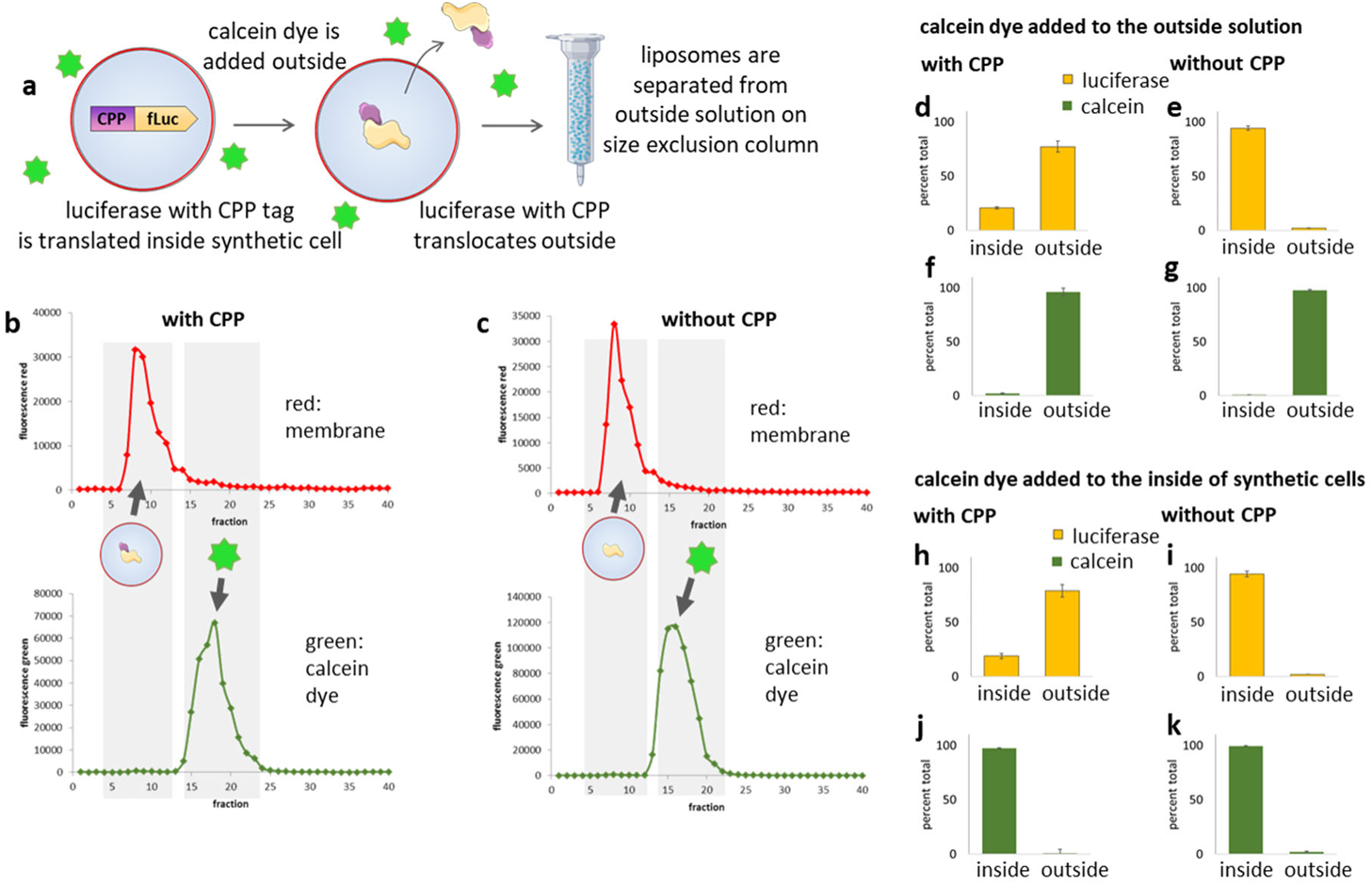
Cell penetrating peptide tag does not increase permeability of synthetic cell membranes. **a**: Schematic of the experiment: synthetic cells expressing protein with or without cell penetrating peptide tags were incubated in the presence of small molecule membrane impermeable green dye calcein. The dye was added to the outside of synthetic cells after liposome formation. Liposome membrane was tagged with red lipid, rhodamine. After 24h of protein expression, samples were purified on a size exclusion column to separate liposome and unencapsulated free solute fractions. **b** and **c**: Example size exclusion purification of synthetic cell containing CPP-tagged luciferase enzyme, after incubation in calcein. The membrane fraction was visualized using red membrane dye (rhodamine). Green channel indicates calcein. **d-k**: percent of luciferase and calcein inside and outside of synthetic cells. **d-g**: Calcein (green bars) was added to the outside of synthetic cells, luciferase (yellow bars) was synthesizes inside cells. **h-k**: Calcein was encapsulated and luciferase synthesized inside synthetic cells. **d**,**f**,**h** and **j**: luciferase with CPP tag. **e**,**g**,**I**,**k**: untagged luciferase. Error bars indicate SEM, n=3. Example size exclusion purification traces are on **Figure S5**. SI figure: luciferase expression by concentration

Molecular communication between populations of synthetic cells is an object of intense engineering efforts.^22^ Having established that we can translocate active enzymes across synthetic cell membrane, we asked whether we can use this system to engineer communication between two populations of synthetic cells.

One population of synthetic cells was designed to express CPP-tagged T7 RNA polymerase (T7 RNAP) under the control of the endogenous *E coli* promoter sigma 70. The other population had a luciferase plasmid under the T7 promoter, but no T7 RNAP was added to those synthetic cells (**Figure 4a**). This resulted in a simple genetic circuit, where T7RNAP from so-called “population T7” was necessary to express luciferase in “population Luc” (**Figure 4b**).

**Figure 4.**
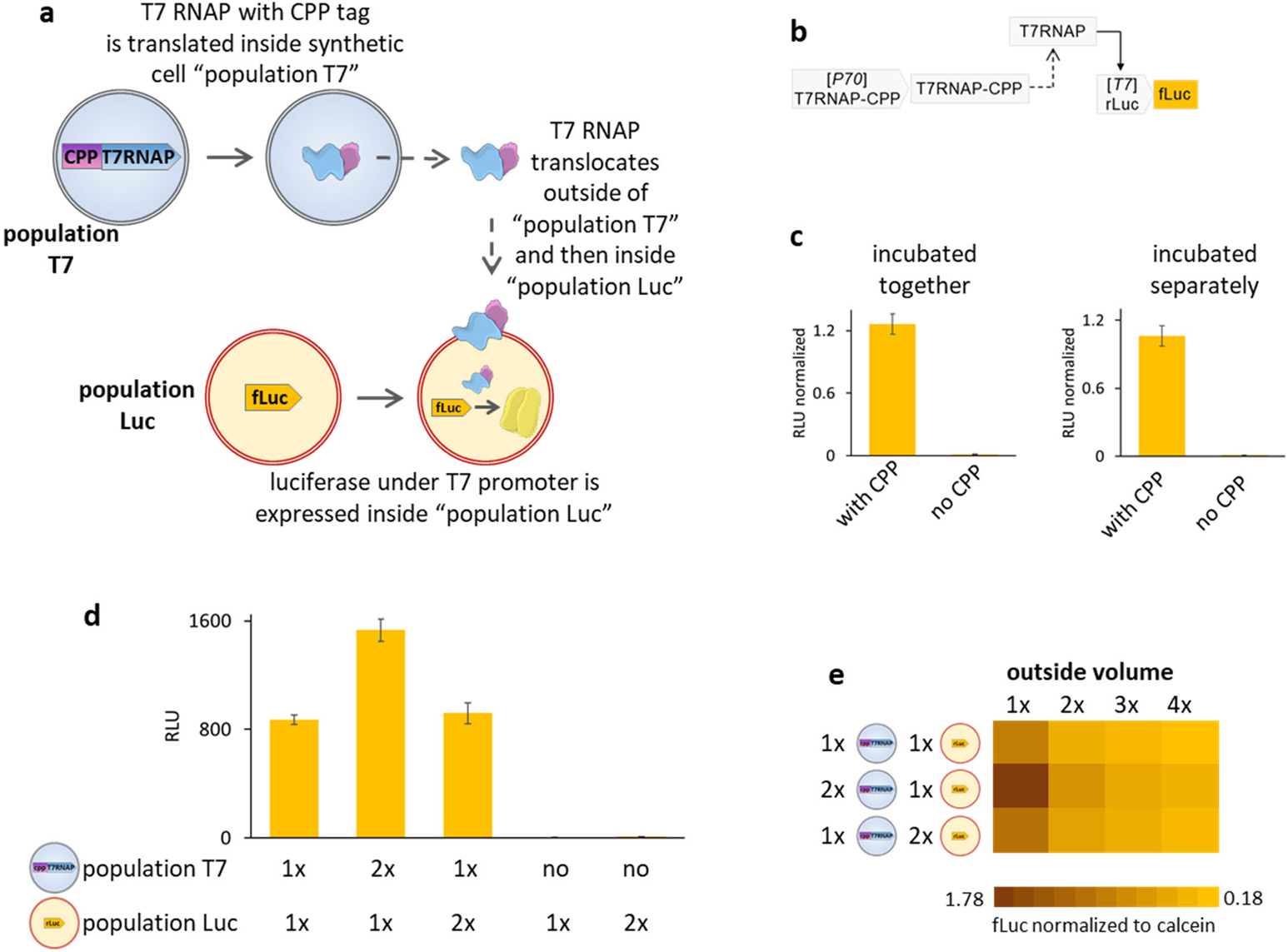
CPP tag facilitates exchange of active enzymes between populations of synthetic cells. **a**: schematic of the experiment: one population of synthetic cells expresses T7 RNA polymerase with a CPP tag. The other population has the luciferase plasmid under a T7 promoter, and red membrane dye. T7 RNAP expressed in “population T7” translocates outside and then inside the “population Luc”, where it initiates synthesis of luciferase. **b:** Schematic of genetic circuit constructed between the two populations. **c:** Luciferase activity in samples with and without a CPP tag on T7RNAP, with “population T7” and “population Luc” either incubated together for 16 hours, or incubated separately for 8h and then combined for another 8h. Luminescence is reported as a ratio between firefly luciferase and control protein renilla luciferase, which was expressed inside the “population T7”. Individual luminescence data for both luciferases are on **Figure S2**. **d:** Luciferase activity measured in samples with different amounts of “population T7” and “population Luc”. The normalized fluorescence data are on **Figure S6**. **e:** Firefly luciferase luminescence normalized to calcein fluorescence (to account for change in the volume of the sample) for samples with increasing volume outside of synthetic cells. 1x is total 10mM lipid concentration, and 2x, 3x and 4x are corresponding dilutions of the whole sample. Individual data points for the heat map are on **Figure S7** and **S8**. Size exclusion purification confirming liposome stability is on **Figure S9**.

When both populations T7 and Luc were incubated together for 16 hours, we observed luciferase activity in samples with CPP-labeled T7 RNAP, but no luciferase was measured in samples with T7 RNAP without a CPP tag (**Figure 4c**). This confirms that the CPP tag is necessary for T7 RNAP to translocate outside of synthetic cells where it is made (the “population T7”) and inside the synthetic cells where luciferase plasmid template is waiting (“population Luc”). Therefore, we established that two populations of synthetic cells can exchange molecular signal, in the form of active RNA polymerase, to complete a genetic circuit.

If the “population T7” is incubated for 8 hours and then mixed with “population Luc” for another 8 hour incubation, the measured luciferase activity is similar to when both populations are incubated for 16 hours together (**Figure 4c**). This suggests that protein expression and equilibration of concentration of T7 RNAP across the membrane occur on similar time scales.

In those experiments, we controlled for total protein synthesis by adding another plasmid to “population T7”: plasmid encoding renilla luciferase. The RLU reported on **Figure 4c** is signal from firefly luciferase normalized to signal from renilla luciferase. Individual luminescence data for both luciferases are shown on **Figure S2**.

To determine whether protein expression or the CPP mediated translocation is the rate limiting factor in this system, we mixed synthetic cells of “population T7” and “population Luc” at different ratios. In samples with twice as much “population T7”, luciferase signal is nearly double the signal from samples mixed at a 1 to 1 ratio. When double the amount of “population Luc” is used, the luciferase signal is similar to the signal measured at 1 to 1 ratio of both populations (**Figure 4d**). This indicates that “population T7” is the rate limiting reagent, since doubling it nearly doubles the measured signal, while doubling the other population does not significantly increase the signal. This is not surprising, as CPP-labeled T7 RNAP needs to not only translocate outside of one synthetic cell population, but it also needs to translocate inside another population.

In absence of either “population T7” or “population Luc” there is no detectable luciferase fluorescence (**Figure 4d**).

Next, we investigated the correlation between the difference in volume inside vs outside of synthetic cells and the efficiency of communication between the two populations.

Synthetic cell populations T7 and Luc were mixed at either equal ratios, or with one of the populations’ concentrations being doubled. Outside volume was also adjusted. 1x is samples where total lipid concentration from 1x “population T7” and 1x “population Luc” is 10mM. Diluting that whole sample with external buffer to 2x, 3x and 4x total volume increases outside volume proportionally. At all outside volume dilutions, the signal followed the previously observed trend: doubling “population T7” significantly increases signal, while doubling “population Luc” has little effect (**Figure 4e**). As outside volume increases, total luciferase signal decreases. This can be explained by the concentration gradient driven CPP mediated translocation. As outside volume increases, the CPP-labeled T7 RNAP that leaves “population T7” is diluted into a larger volume, thus decreasing the relative concentration gradient between the outside and inside of “population Luc”, thus slowing down translocation inside “population Luc” (**Figure 4e**).

Cell penetrating peptides, and penetratin in particular, have been demonstrated to carry cargo inside live cells.^23^ We asked whether our penetratin tag can carry nucleic acid cargo across a synthetic cell membrane.

Synthetic cells were designed with a CPP-labeled MS2 RNA binding domain, which is a viral coat protein binding specific RNA motif with very high affinity.^24,25^ Inside those cells, we also placed the fluorescent RNA aptamer broccoli with an MS2 RNA domain added at the 3’ end. The CPP-tagged MS2 protein can bind broccoli and translocate outside of synthetic cells (**Figure 5a**). After a 16 hour incubation, samples were purified on a size exclusion column, and broccoli fluorescence was measured inside and outside synthetic cells.

**Figure 5.**
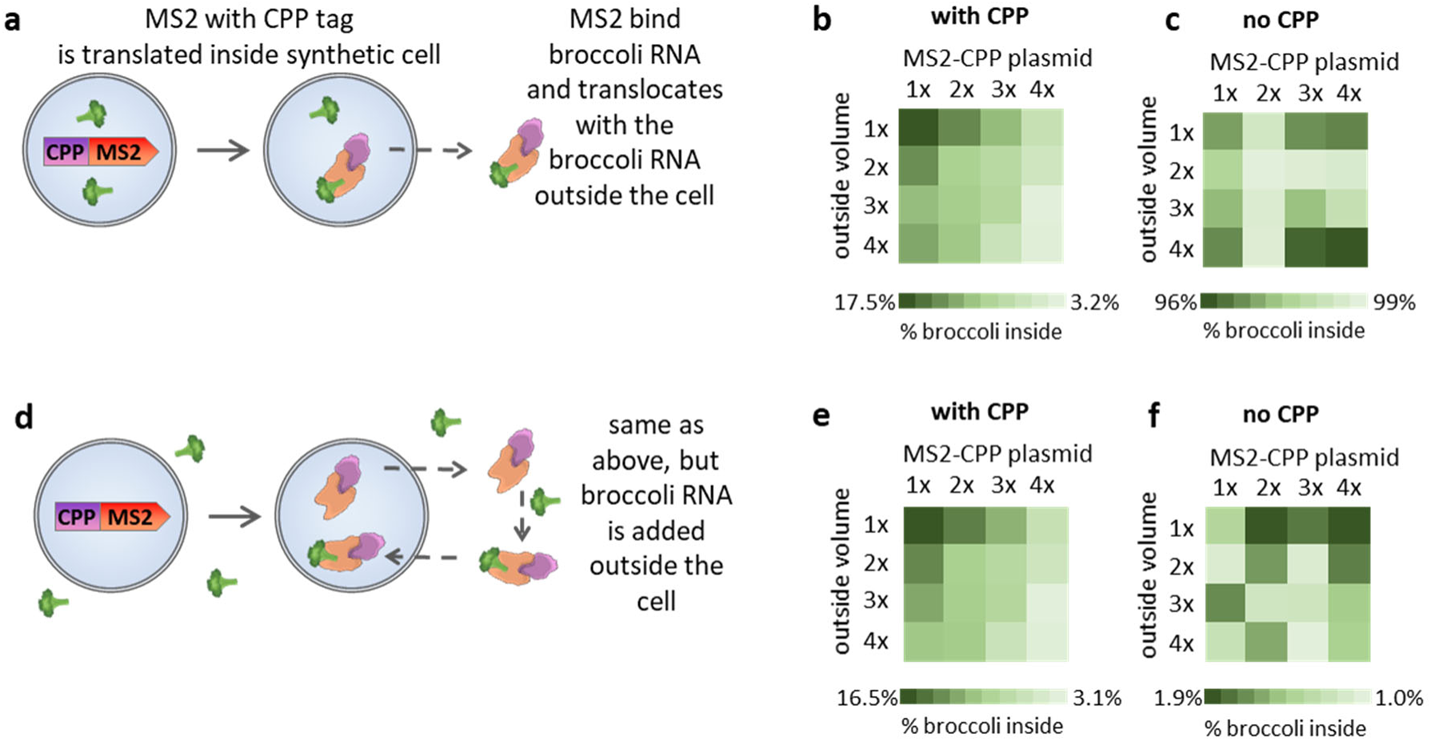
CPP tag facilitates export and import of RNA payload. **a: Export of RNA**: Synthetic cells are filled with broccoli fluorescent RNA aptamer with MS2 RNA domain. The cells express CPP-tagged MS2 RNA binding domain. CPP-MS2 protein binds broccoli-MS2 RNA and translocates outside of the cell. **d: Import of RNA**: Experiment set up same as on panel **a**, but the broccoli fluorescent RNA aptamer with MS2 RNA domain is added outside of synthetic cells, and transported inside the cell by the CPP-tagged MS2 RNA binding domain. **b**, **c**, **e** and **f**: The % of fluorescent RNA broccoli inside synthetic cells after 16h reaction and size exclusion purification. Outside volume 1x is sample with 10mM lipid. 2x, 3x and 4x are corresponding dilutions of the whole sample. Plasmid 1x is 2nM. 2x, 3x and 4x are corresponding increased plasmid concentrations. All individual data points are on **Figure S10** and **Figure S11**. Example size exclusion purification traces are on **Figure S12**. **b** and **c**: Export of RNA. **e** and **f**: Import of RNA. **b** and **e**: Samples with CPP tag on MS2 protein, **c** and **f**: no CPP tag on MS2 protein. Import and export experiments were also performed with translation system and CPP-MS2 plasmid outside of synthetic cells, results on Figures S13 and S14.

The amount of broccoli aptamer remaining inside synthetic cells after incubation and purification strongly correlates with both the amount of CPP-MS2 protein expressed inside the cell and the outside volume (**Figure 5b**). The 1x MS2-CPP plasmid indicates samples with 2nM plasmid, and 2x, 3x and 4x samples indicate proportional increase in plasmid concentration. As the amount of CPP-MS2 plasmid increases, the amount of broccoli remaining inside cells decreases. This indicates that the availability of the CPP-MS2 protein to shuttle broccoli across the membrane was the rate limiting factor in the translocation efficiency. Another factor increasing translocation yield is outside volume, as expected for concentration driven process. 1x outside volume corresponds to samples with 5mM total lipid and 2x, 3x, and 4x samples are corresponding dilutions, increasing total sample volume but not internal synthetic cell volume. As outside volume increases, less broccoli remains inside synthetic cells (**Figure 5b**).

In samples without CPP, neither the amount of MS2 plasmid nor the outside volume correlate with the amount of broccoli inside, and most of the broccoli aptamer remains inside the synthetic cells under all conditions (**Figure 5c**).

We also investigated an opposite scenario: broccoli added to the outside of a synthetic cell, while the CPP-MS2 protein is expressed inside (**Figure 5d**). The amount of broccoli inside the synthetic cell after a 16 hour incubation and purification follows the same trends as previously observed: more CPP-MS2 plasmid and a larger outside volume results in more broccoli outside the cell (**Figure 5e**). In samples without a CPP tag on MS2, most of the broccoli remains outside the cells under all tested conditions (**Figure 5f**).

It is interesting to note that the final amount of broccoli inside synthetic cells is similar after a 16 hour incubation in samples with a CPP tag on MS2 whether broccoli started out inside (**Figure 5b**) or outside synthetic cells (**Figure 5e**). This indicates that a 16 hour incubation time is enough to reach steady state of translocation in and outside the cell.

The membrane transport is non-directional – the CPP tag can translocate the same protein both ways across the membrane, in a process that is largely concentration driven. We hypothesized that we could improve accumulation of cargo on one side of the membrane by removing the CPP tag from the cargo after transport. To test it, we used a coiled-coil split intein system: the coiled-coil dimerizing proteins bring two halves of the split intein together, and the intein can irreversibly splice.^26,27^ We created a construct where the CPP tag was attached to the intein tag (INT) attached to firefly luciferase (**Figure 6a**). The other half of INT tag was added to the outside of synthetic cells as purified protein. After the CPP-INT tagged luciferase is expressed and translocates outside of synthetic cells, the other half of INT tag dimerizes and splices, removing the CPP tag from luciferase. This prevents luciferase from diffusing back into the synthetic cell. The luciferase export can therefore be either irreversible (INT tag splices to remove CPP tag), reversible (no INT tag, so CPP tag cannot be spliced out), or in control samples no export at all (no CPP tag). We also tested influence of external volume – 1x is sample with 5mM lipid; 2x, 3x and 4x are respective dilutions of the 1x sample, increasing the volume outside synthetic cells without changing the internal volume.

**Figure 6.**
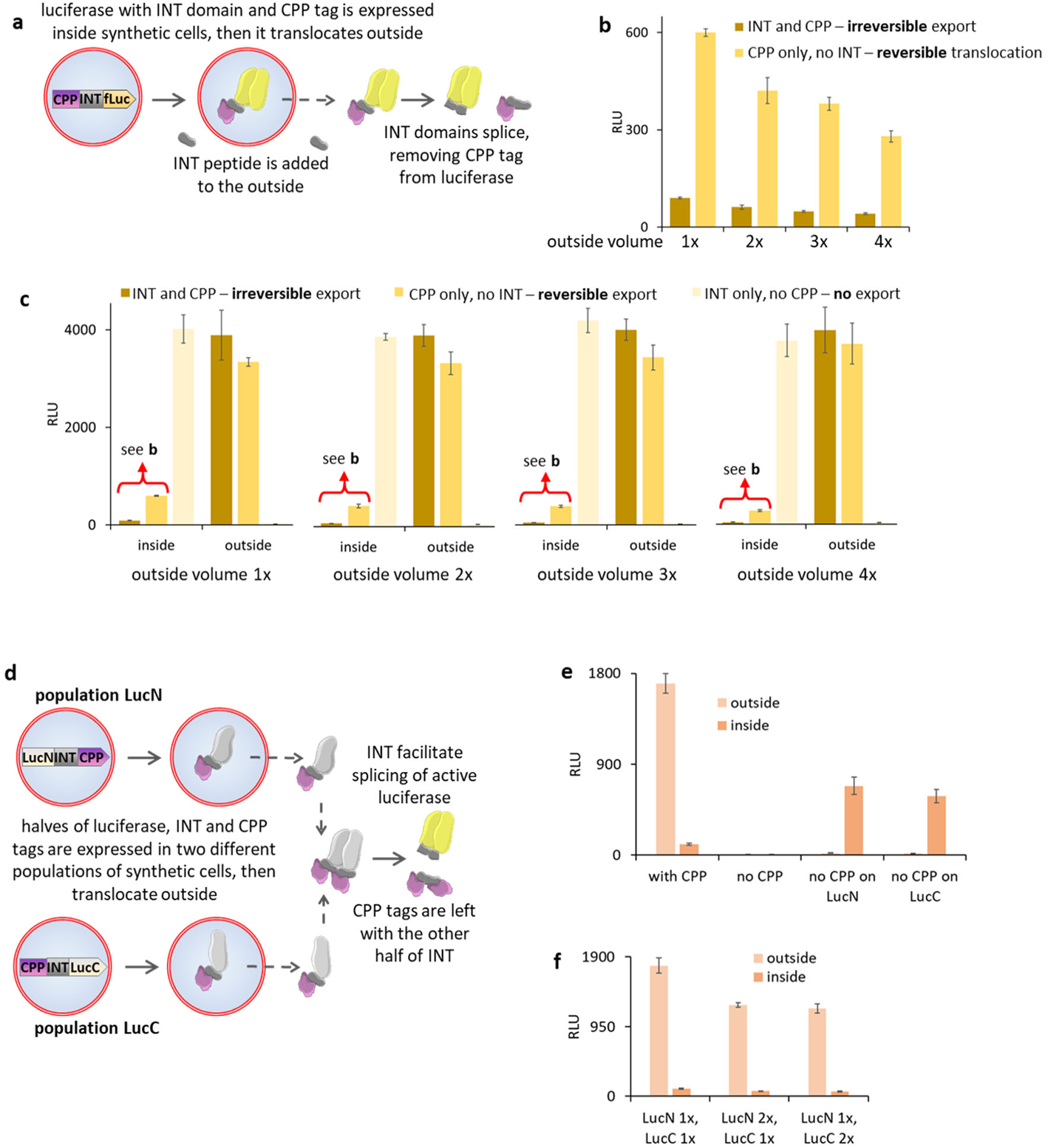
Export and import of proteins can be controlled by dynamically removing the CPP tag. **a**: Synthetic cells express CPP tagged luciferase with a coiled-coil intein (INT) domain. The other half of split intein domain is added outside synthetic cells. After export of CPP tagged luciferase, the inteins splice, removing CPP tag from luciferase and trapping the luciferase outside of synthetic cell. **b** and **c:** Luminescence of luciferase inside and outside of synthetic cells after 16h reaction and size exclusion purification. Samples with irreversible export (INT and CPP, the CPP tag is spliced out after export), reversible export (CPP only, no INT, so CPP tag cannot be spliced out), and without export (INT only tag, no CPP tag). The red curly brackets highlight data for luciferase inside synthetic cells with irreversible and reversible export, those data points are shown in higher resolution on panel **b**. Outside volume 1x is sample with 10mM lipid. 2x, 3x and 4x are corresponding dilutions of the whole sample. Error bars indicate S.E.M., n=3. **d**: Two population of synthetic cells express two different halves of split luciferase, each half with CPP tag and INT splicing tag. The halves translocate across the synthetic cell membrane and INT splice, separating CPP tag from luciferase. **e:** Luminescence of luciferase inside and outside of synthetic cells after 16h reaction and size exclusion purification. Samples with CPP on both halves of luciferase, without any CPP tag, and with CPP tag only on either LucN or LucC half. Error bars indicate S.E.M., n=3. **f**: Luminescence of luciferase inside and outside of synthetic cells after 16h reaction and size exclusion purification. Populations expressing LucC and LucN were mixed at different ratios, with final total lipid concentration always the same (5mM total lipid). Error bars indicate S.E.M., n=3.

After a 16 hour reaction and size exclusion purification, we quantified luciferase activity inside and outside of synthetic cells (**Figure 6c**). Samples with reversible export had significantly more luciferase inside synthetic cells than samples with irreversible export. This indicates that the irreversible export scheme allowed for accumulation of the product outside synthetic cells at larger concentrations than simple concentration driven equilibration in reversible export samples. The difference in luciferase export efficiency for reversible export samples followed the expected outside volume dilution – more luciferase was exported if outside volume was larger. This is consistent with previous observations for GFP (**Figure 1j**), T7 RNAP (**Figure 4d and e**) and for MS2 RNA payload (**Figure 5b and e**).

However, for the case of irreversible export, the difference in the amount of luciferase exported at different outside volumes was much less significant (**Figure 6b**). This could be explained by the shifting of concentration equilibrium with each irreversible cleavage of CPP tag. The luciferase freed from the CPP tag was removed from the pool of translocation-competent proteins, leaving smaller CP tagged pool to equilibrate.

Inside and outside luciferase activity data for all samples is on **Figure 6c**, with data points for luciferase inside synthetic cells with irreversible and reversible export repeated on **Figure 6b** for better visibility. As expected, in all cases of samples without a CPP tag, there was no luciferase activity outside of synthetic cells.

Encouraged by the results from the cleavable INT tag, we designed another iteration of the intracellular molecular signal exchange scheme. This time, two populations of synthetic cells expressed two halves of a split luciferase, each attached to a CPP-INT tag.^26^ The split luciferase is not active until both halves are properly joined.^28^ When the two halves meet, the INT tags splice, removing CPP tags from luciferase and restoring luciferase activity (**Figure 6d**). This design takes advantage of irreversible export scheme, and only spliced luciferase (unable to cross the membranes anymore) is detectable in activity assay. The results indicate that most of the splicing occurs outside of both populations of synthetic cells, as most of the luciferase activity was found in the fraction outside (**Figure 6e**).

Interestingly, we found small but measurable luciferase activity inside synthetic cells as well. Since we found no luciferase activity at all in samples where neither half had a CPP tag, we know that luciferase without a CPP tag cannot cross the membrane. The active luciferase found inside synthetic cells can be explained by the location of the splicing event. In most cases, N and C terminal luciferase halves exit the respective populations where they were synthesized, encounter their splicing partner, and remain outside after splicing (as the CPP tag is removed in splicing). In a small amount of cases, one of the halves of luciferase must have translocated twice: outside of its population of origin and inside the other population, where it encountered a splicing partner, and resulting active luciferase was stuck inside the cell. (**Figure 6e**).

Both halves of luciferase equally contributed to the final signa, indicating no preferential translocation of one or the other protein: the highest signal was observed when both populations of synthetic cells were mixed in equal proportions (**Figure 6f**).

To test the efficiency of the scheme requiring a single half to translocate twice (outside of population or origin and inside the other population), we did experiments where only one half of the luciferase had a CPP tag. The only way that luciferase activity could be restored was if the CPP tagged half enters the cell where the half with the INT but no CPP was waiting. The resulting active luciferase would then be stuck inside the synthetic cell (since CPP tag is excised in splicing). In both cases (CPP tag on either N or C half), we detected luciferase activity inside synthetic cells, and as expected no activity was found outside. (**Figure 6e**).

Those experiments demonstrate that it’s possible to drive export and import of CPP tagged cargo across synthetic cell membranes and between different synthetic cell populations beyond the simple concentration gradient.

To expand the cell-to-cell communication capacities, we engineered a molecular communication system based on mRNA payloads. One population of synthetic cells (population MS2) had a CPP-tagged MS2 RNA binding domain and a plasmid for the small peptide FlAsH^29^. The FlAsH peptide binds an arsenic ligand that becomes fluorescent upon binding to the FlAsH peptide. In this synthetic cell population, both CPP-tagged MS2 and FlAsH peptide are being translated, but because of the lack of the arsenic ligand, the FlAsH peptide is not fluorescent. The FlAsH mRNA was tagged with an MS2 motif in its 3’ UTR. Another population of synthetic cells (population FlAsH) contained all components of TXTL system and the FlAsH ligand, but no DNA template or RNA polymerase. The CPP tagged MS2 protein from “population MS2” binds to the MS2 tag on the FlAsH mRNA, translocating outside the cell and inside the other “population FlAsH.” Inside “population FlAsH,” the FlAsH mRNA is translated, and presence of the FlAsH ligand enables fluorescence (**Figure 7a**).

**Figure 7.**
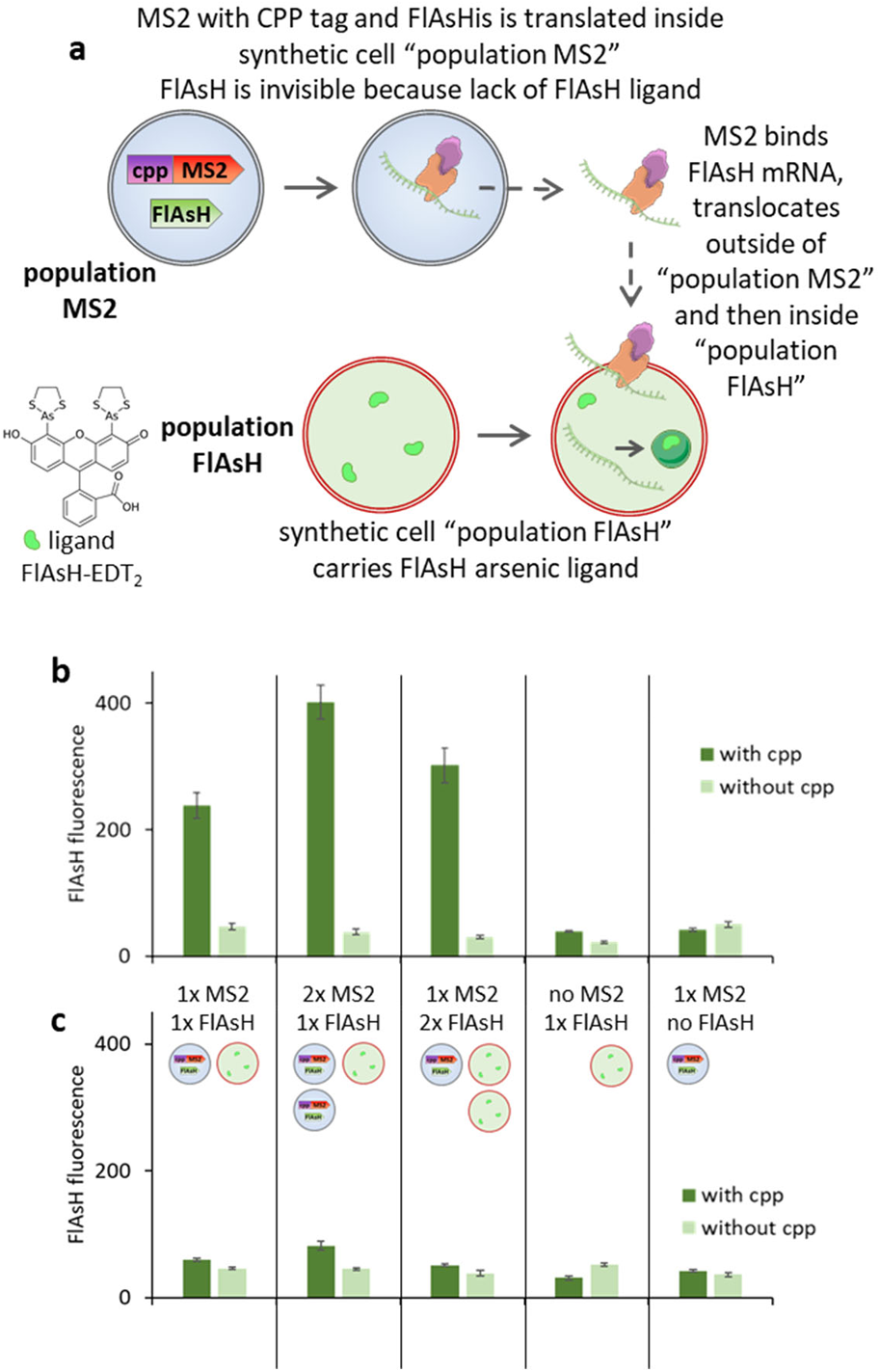
Exchange of mRNA signals between populations of synthetic cells. **a**: Schematic of the experiment. One population of synthetic cells expresses CPP tagged MS2 RNA binding protein, and FlAsH mRNA with MS2 tag. The CPP tagged MS2 protein binds MS2 domain on FlAsH mRNA and translocates across the membrane. Second population of synthetic cells contains no DNA templates, but it contains ligand for FlAsH peptide. The CPP MS2 protein with FlAsH mRNA payload translocates inside the second population. FlAsH is expressed from mRNA, and FlAsH protein fluorescence is detectable thanks to presence of FlAsH ligand. **b**: Fluorescence of FlAsH protein inside synthetic cells, for samples with and without CPP tag on MS2 protein. Results for samples without MS2 tag on FlAsH mRNA are on **figure S15**. **c**: Experiment same as on panel **b**, with RNAse A added outside of synthetic cells. Control experiments for RNAse A membrane permeability are on **figure S16**. Experiments performed with different ratio of population MS2 and population FlAsH. 1x equals liposomes with 5mM lipid, 2x is 10mM lipid. Fluorescence measured after 16h incubation. Error bars indicate S.E.M., n=3.

We observed FlAsH peptide fluorescence in samples with both populations of synthetic cells mixed together (**Figure 7b**). Mixing twice the amount of population MS2 (10mM lipid vs 5mM) we observed nearly double the FlAsH fluorescence, while doubling “population FlAsH” produced only a minor increase in signal. In light of other observations presented in this paper, this is not surprising. The data suggests that the rate limiting step in those reactions is the availability of the mRNA translocated across two synthetic cell membranes.

Over the course of those experiments, we discovered a practical limit on payload size in this system. We were unable to translocate larger RNA, despite testing several constructs prepared with varying lengths of mRNA capped with the MS2 domain. We were unable to observe any RNA transport for payloads larger than the FlAsH mRNA.

To enable engineering of another control point in the communication system, we targeted the translocating mRNA for degradation by adding RNAse A to the outside of synthetic cells. The FlAsH fluorescence decreased significantly, with only a very small recovery in the 2x “population MS2” experiment (**Figure 7c**). This indicates that most of the FlAsH mRNA is digested by the extracellular RNAse. This provides another means of controlling molecular communication between synthetic cells, enabling interception and inactivation of the mRNA signal as its being transported between the cells.

The membrane translocation system we demonstrated here opens up new possibilities for synthetic cell engineering, bioproduction and “green chemistry” engineering. While a lot of specific applications remain to be explored, here we demonstrate the chassis for engineering synthetic cells that can export protein products and nucleic acid payloads as well as demonstrate a new way of engineering communication between different populations of synthetic cells.

We demonstrated that the CPP-mediated membrane transport is driven by the concentration difference, and the transport can be pushed to higher product accumulation by removing the CPP tag. We also demonstrated that the CPP tag does not increase permeability of the synthetic cell membrane. We engineered a new mode of protein and nucleic acid driven molecular communication between different populations of synthetic cells, and we demonstrated that nucleic acid mediated communication can be modulated by use of nucleases. The nucleic acid mediated communication is of special interest for engineering complex behaviors of synthetic cell populations, as the molecular message can be encoded in infinitely programmable nucleic acid payload.

These results establish the possibility to create highly controlled minimal cells with the capability to export selected cargoes. In addition to advancing the field of synthetic cell engineering towards construction of a live synthetic minimal cell, the technology demonstrated here might prove to be important in areas such as drug delivery, diagnostics, metabolic engineering and environmental remediation.

## Acknowledgments

We thank Dr. Emmanouil Karagiannis for helpful discussions about cell penetrating proteins that inspired this project. We thank Dr. Vincent Noireaux for discussion about protein expression and cell-free translation systems, and for the gift of T7 RNA polymerase plasmid. This work was supported by National Aeronautics and Space Administration award 80NSSC18K1139, National Science Foundation. awards 1844313 and 1807461, Semiconductor Research Corporation grant number 2018-SB-2837-C and NIH award 5R01MH114031.

## Author contributions

JMH, KS, NJG, CD, JG-G, BC, MRP and KPA performed experiments. All authors analyzed data. JMH, NJG, AEE and KPA wrote the paper.

## Materials and methods

Fluorescence and luminescence were measured using Molecular Devices SpectraMax plate readers. All measurements of fluorescence for samples directly compared with each other (on the same figure panel, or on associated panels) were done using the same PMT settings.

### Preparation of POPC: POPG:cholesterol liposomes

POPC (1-Palmitoyl-2-oleoyl-sn-glycero-3-phosphocholine), POPG (1-palmitoyl-2-oleoyl-sn-glycero-3-phospho-1’-rac-glycerol) and cholesterol were purchased from Avanti Polar Lipids. Stock solutions of 10mg/mL in chloroform were used for making lipid films. Liposomes were prepared according to previously described protocol^30^. Unless otherwise stated, we used liposomes made of 10mol% POPG 90mol% POPC with 200mol% cholesterol relative to total lipid concentration. Liposomes were most commonly prepared with 5mM total lipid. Aliquots of lipids were mixed and solvent was removed by overnight evaporation. Lipid samples were suspended in mineral oil and incubated, with vigorous shaking, overnight. Aqueous phase containing TXTL components, plasmids and other molecules was added to the mineral oil suspension with vigorous mixing. Emulsion samples were pipetted on top of centrifugation buffer (100 mM HEPES + 200 mM glucose, pH 8) and centrifuged at 18000 rcf at 4°C for 15 minutes. Top oil phase was removed, and buffer containing liposomes was moved to fresh tube on top of equal volume of wash buffer (100 mM HEPES + 250 mM glucose, pH 8). Samples were centrifuged at 12000 rcf at 4°C for 5 minutes. Liposome containing buffer was collected from the bottom of the tube.

### TXTL protocol

The E coli TXTL was prepared according to the previously described protocol.^31^ Each reaction contained 1.5mM ATP and GTP, 0.9mM CTP and UTP, 0.068 mM folinic acid, 200 ug/mL of E. coli tRNA mixture, 0.33 mM nicotinamide adenine dinucleotide (NAD), 0.26 mM coenzyme-A (CoA), 1.5 mM spermidine, 4 mM sodium oxalate, 0.75mM cAMP, 30mM 3-PGA, 50mM HEPES pH 8, 1mM DTT, 12mM Mg-glutamate, 140mM K-glutamate, 2mM of each of the 20 amino acids, 1x RNAse inhibitor Murine (NEB).

For complete liposome reaction, 6ul Mg-glutamate, 23.33ul K-glutamate, 50ul energy mix, 50ul amino acids, 7ul 10mM DTT, 8.54ul 207nM plasmid template, 1.4ul RNAse inhibitor, 23.33ul cell extract and 2.76ul 50uM T7 were mixed. with water added to 70ul.

All TXTL reactions were incubated at 30°C for 16 hours, unless stated otherwise.

To increase reproducibility and comparability of data, all experiments directly compared to each other (on the same figure panel, or on associated panels) were done using the same batch of cell extract.

### Size exclusion purification of synthetic cell liposomes

Sepharose 4B (Sigma Aldrich) was poured into gravity flow column (BioRad). Sepharose was washed with minimum 4 volumes of running buffer. For most synthetic cell purifications, 50mM HEPES pH 8 was used to match the buffer of TXTL reaction. Sample of liposomes was carefully loaded on top of the Sepharose column, and eluted from the column using 3 to 6 volumes of running buffer. Fractions were collected into 96 well plate using fraction collector (Gilson). Fluorescence was red using plate reader directly from the collection plate to determine placement of the liposome and free dye fractions. If needed, liposome and free dye fractions were collected from the well plate and pooled together.

### Luciferase luminescence assays

Before each assay, liposomes were lysed using 0.1% v/v of Triton. Firefly luciferase activity was measured using Steady-Glo luciferase assay system (Promega). For measuring firefly and renilla luciferase activity in the same sample, Dual-Glo assay (Promega) was used. Both kits were used according to manufacturer’s instructions.

### Western Blot protein expression analysis

Samples were analyzed using 7.5% SDS-Page resolving gel. Most gels were run at constant 100V for 1h. Transfer buffer was 25mM Tris, 192mM Glycine, 10% methanol. PVDF membranes were soaked in methanol until wet, then soaked in transfer buffer. Transfer was performed at 100v for 1 hour at room temperature. After transfer, membrane was moved into blot contained and blocked with 10 mL of 5% milk powder in TBST for 1 hour at room temperature. Membrane was then covered with solution of primary antibody and incubated at 4°C overnight with shaking. Primary antibody was removed and membrane was washed three times with TBST buffer (20mM Tris, 150mM NaCl, 0.05% tween) for 10 minutes each wash. Secondary antibody solution was added and incubated for 1 hour. The secondary antibody solution was removed, membrane was washed three times with TBST for 10 minutes each wash. A 1:1 mixture of Stable Peroxide Solution and Luminol Enhancer Solution was added on top of the membrane and the membrane was visualized after 5 minutes of incubation.

## Supplementary Information

### Liposome volume calculation

Below is the rationale for calculation of liposome volume (lumen as % of total sample volume). This calculation has been originally published in our earlier work^27^, and the interactive calculator using those formulas is available on our lab website: http://protobiology.org/tools-liposome.php

%internal volume = vol_liposome*liposomes_ml*10-19

Where:

vol_liposome = (4/3)*Pi*(r3)

is the volume of the lumen of a single liposome, in nm3;

liposomes_ml = surface_area_ml/area_liposome is the number of liposomes per 1 mL; surface_area_ml = (c*10^-6)*((760*10^21)/0.9*NA)/2.5)/2

is the surface area of liposomes per 1 mL of solution of a given c (mM), with POPC MW=760 and length of the lipid bilayer approximated to 2.5nm; NA is Avogadro’s number;

and finally,

area_liposome = 4*Pi*(r2)

is the surface area of the liposome outer leaflet, in nm2.

These calculations were made with the assumption that liposome curvature is negligible, so the inner and outer leaflet contain an equal number of lipids and have equal surface area. The thickness of the bilayer was approximated at 2.5 nm^32^.

The addition of cholesterol increases bilayer thickness up to 30%, thus affecting the encapsulation rate^33^, but we cannot reliably estimate the influence of cholesterol on packing density and surface area of the liposomes.

According to this formula, a 25 mM solution of 200 nm POPC liposomes will contain ∼14% of the total volume encapsulated inside liposomes.

In reality, the encapsulation rate of liposomes used in our experiments is likely lower. This is due to factors like the presence of cholesterol in POPC membranes and the fact, that in liposomes extruded through, e.g., a 200 nm filter, the size distribution of liposomes varies greatly and is, on average, smaller than 200nm^34–36^.

The differences in yield of protein synthesis inside synthetic cells, explained by the difference in efficiency of encapsulating the TXTL enzyme mix, have been observed^37^.

---

**Figure S1.**
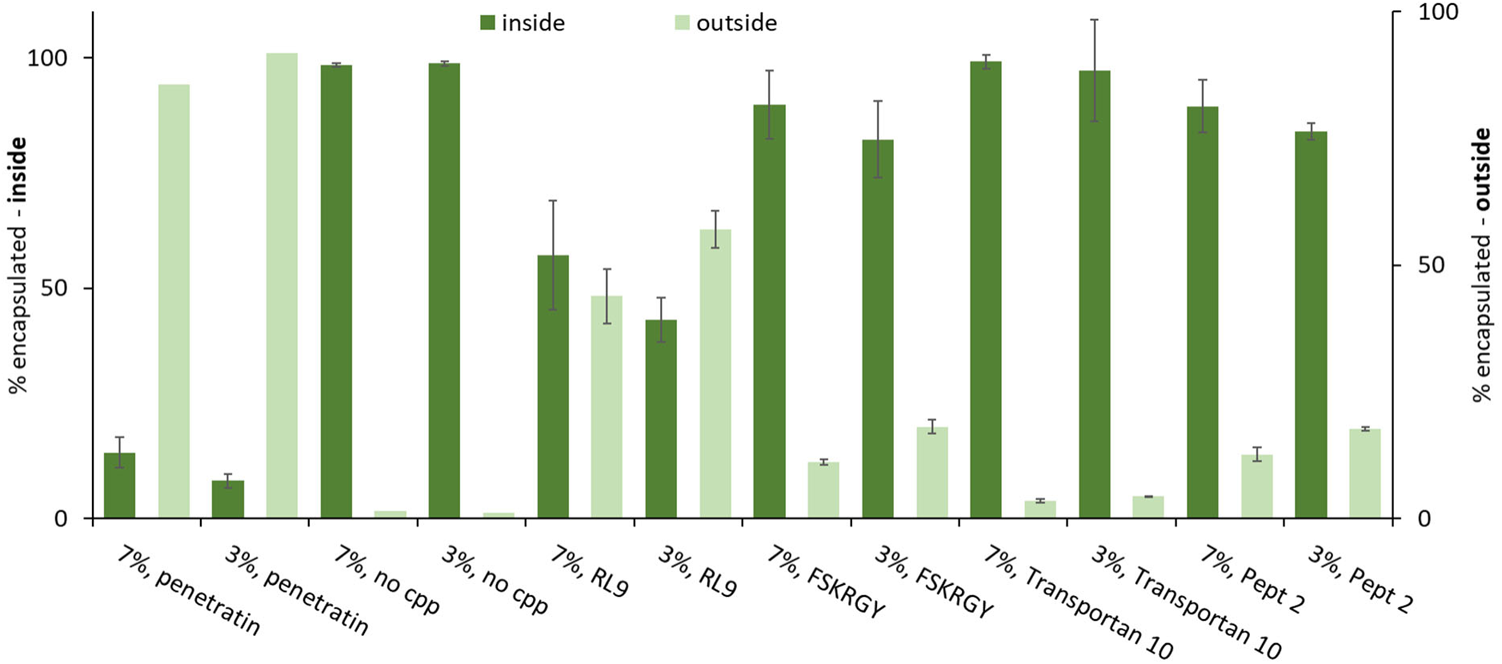
Initial testing of different cell-penetrating peptides. GFP was labeled with CPP sequence. Synthetic cells expressing GFP were purified on size exclusion column after 18h reaction following the protocol used in **Figure 1j** and **1k**. 7% and 3.5% corresponds to the percentage of the whole sample that is encapsulated inside the lumen of synthetic cells. Error bars indicate S.E.M., n=3.

**Figure S2.**
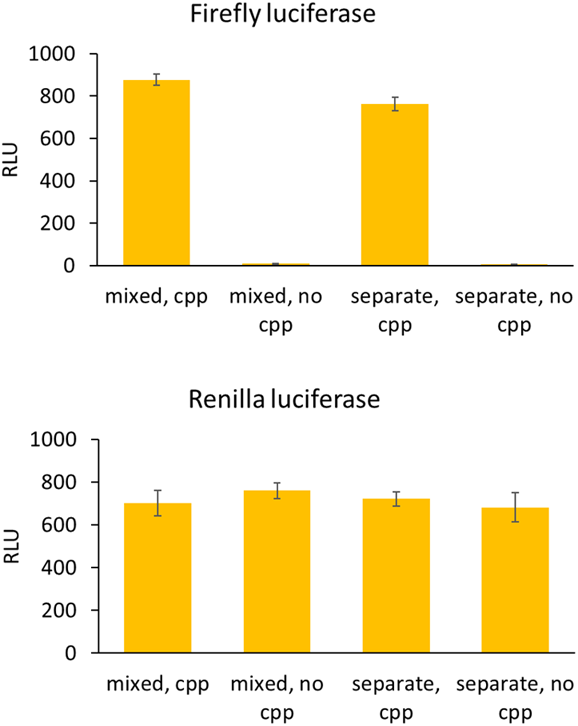
Individual sample firefly luciferase and renilla luciferase values for incubation test experiments shown on figure 4c. Error bars indicate SEM, n=3.

**Figure S3.**
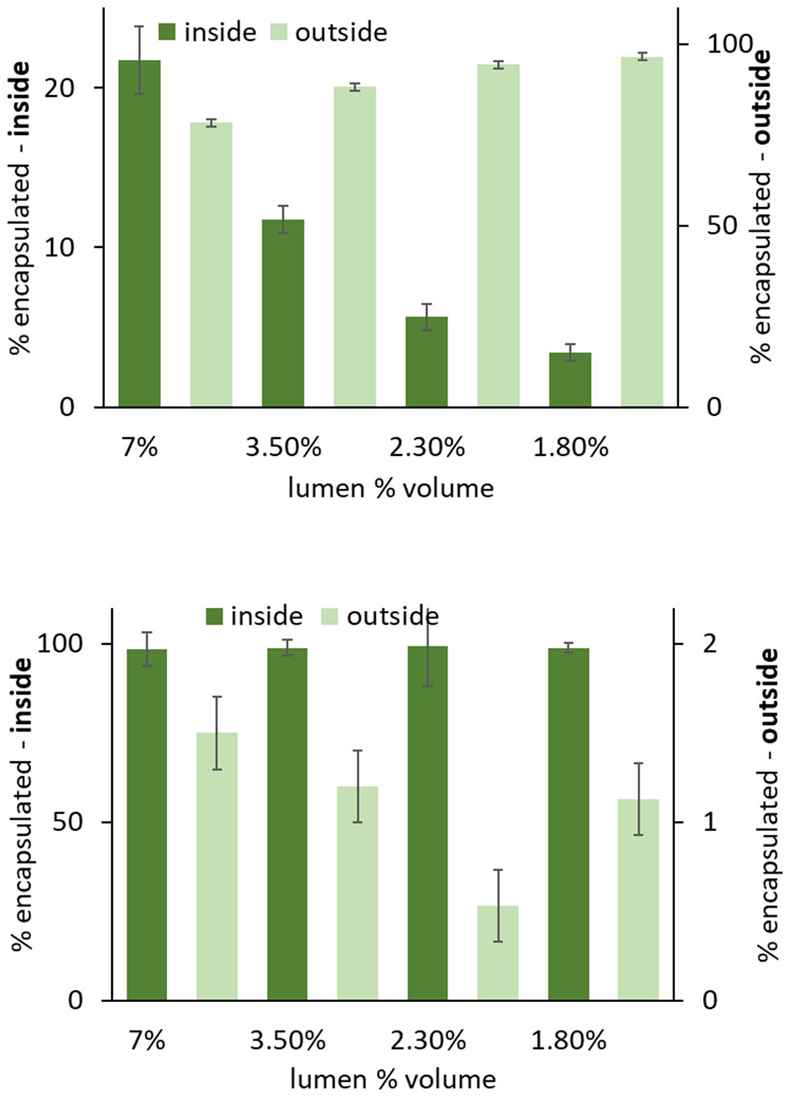
Percent of GFP encapsulated inside (left Y axis) and percent of GFP outside the cell (right Y axis) for samples containing different amount of synthetic cells, corresponding to different % of the total sample volume being encapsulated inside the vesicles. Experiment setup is identical as for data shown on Figure 1j and k, but GFP plasmid concentration is 1nM. See **Table yyS1** for liposome concentrations corresponding to different lumen volume. Error bars are S.E.M, n=3.

**Figure S4.**
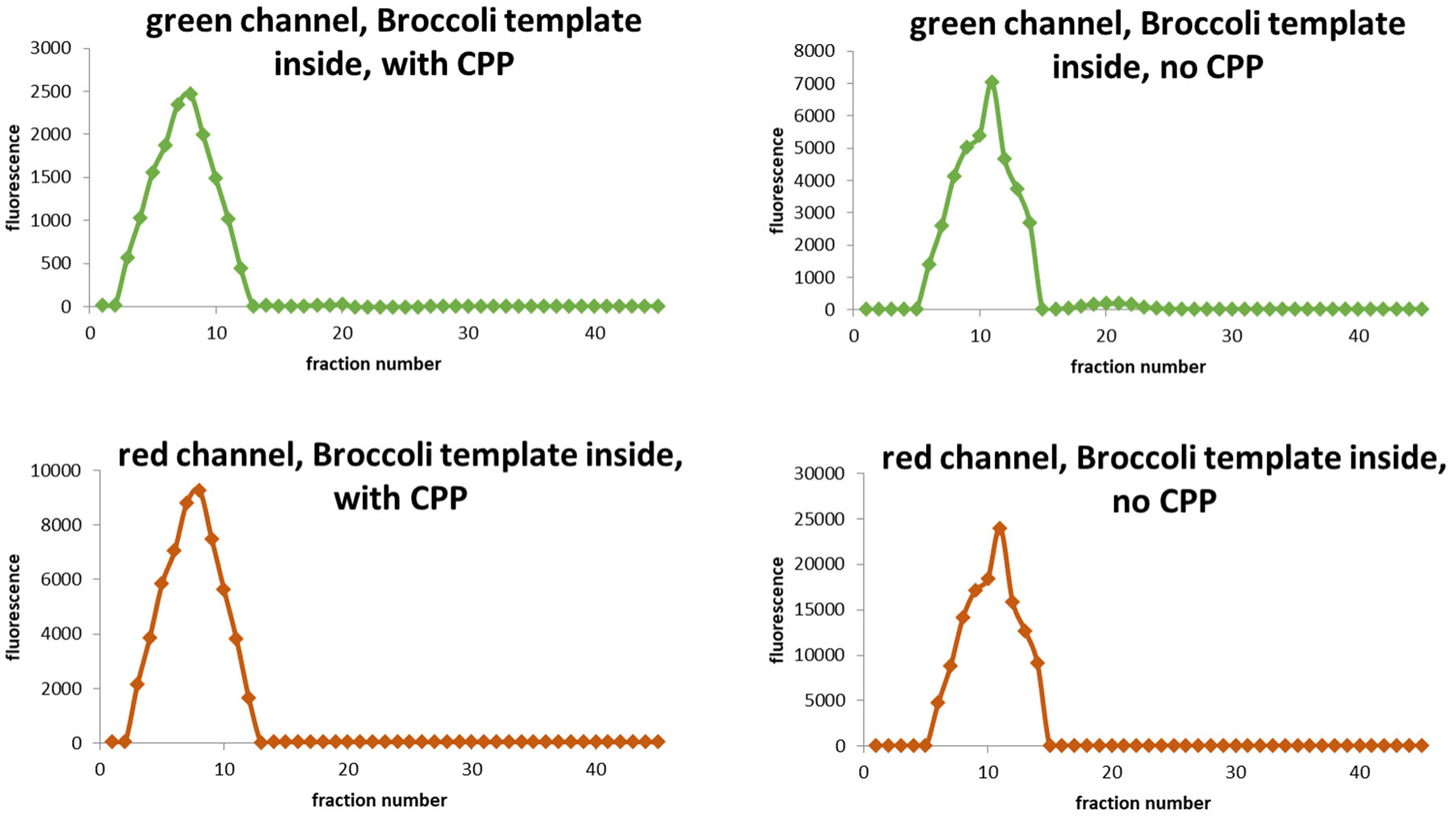
Size exclusion purification data. Synthetic cell liposomes were purified on Sepharose 4B. Fluorescence was measured in green channel (Broccoli) and red channel (Rhodamine membrane dye).

**Figure S5.**
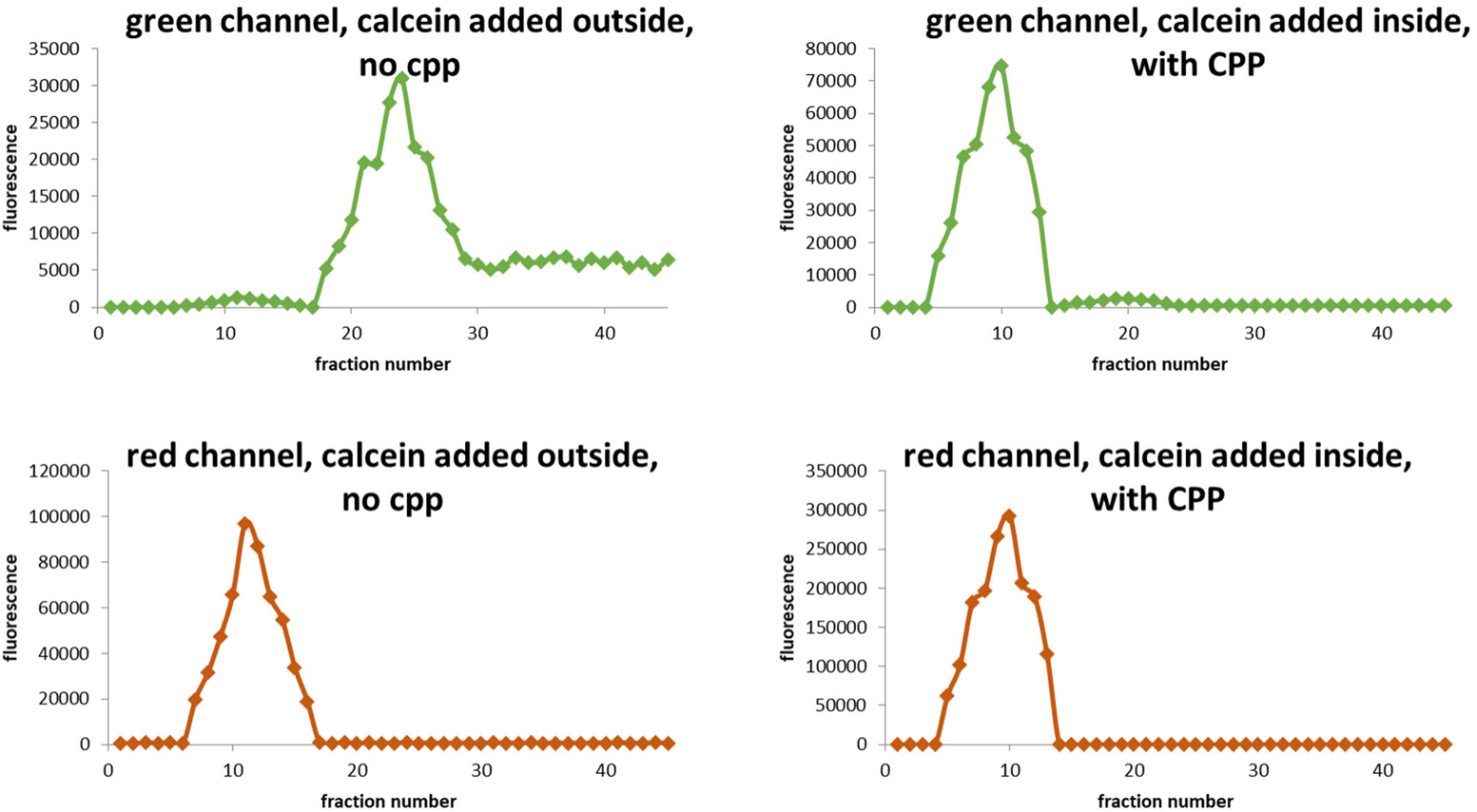
Size exclusion purification data. Synthetic cell liposomes were purified on Sepharose 4B. Fluorescence was measured in green channel (calcein) and red channel (Rhodamine membrane dye).

**Figure S6.**
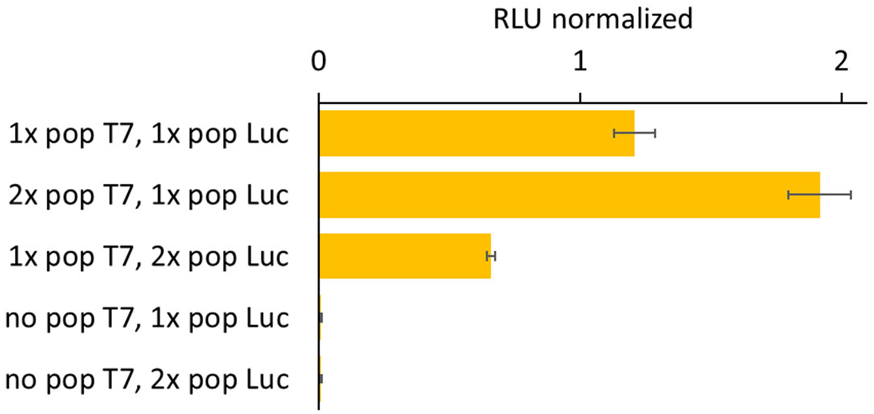
Fluorescence data normalized (Firefly / Renilla signal) for samples shown on figure 4d. Error bars indicate SEM, n=3.

**Figure S7.**
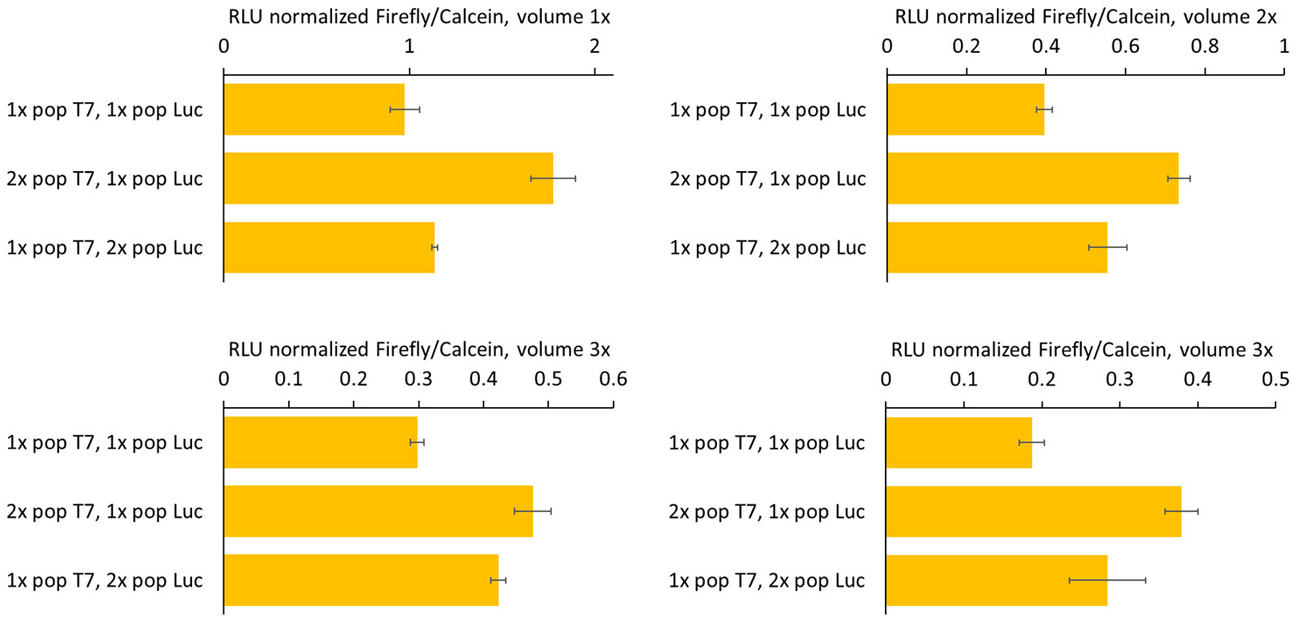
Individual experiment data for the heatmap showed on Figure 4e, Firefly luciferase luminescence normalized to calcein fluorescence to normalize for volume of the sample. Error bars indicate SEM, n=3.

**Figure S8.**
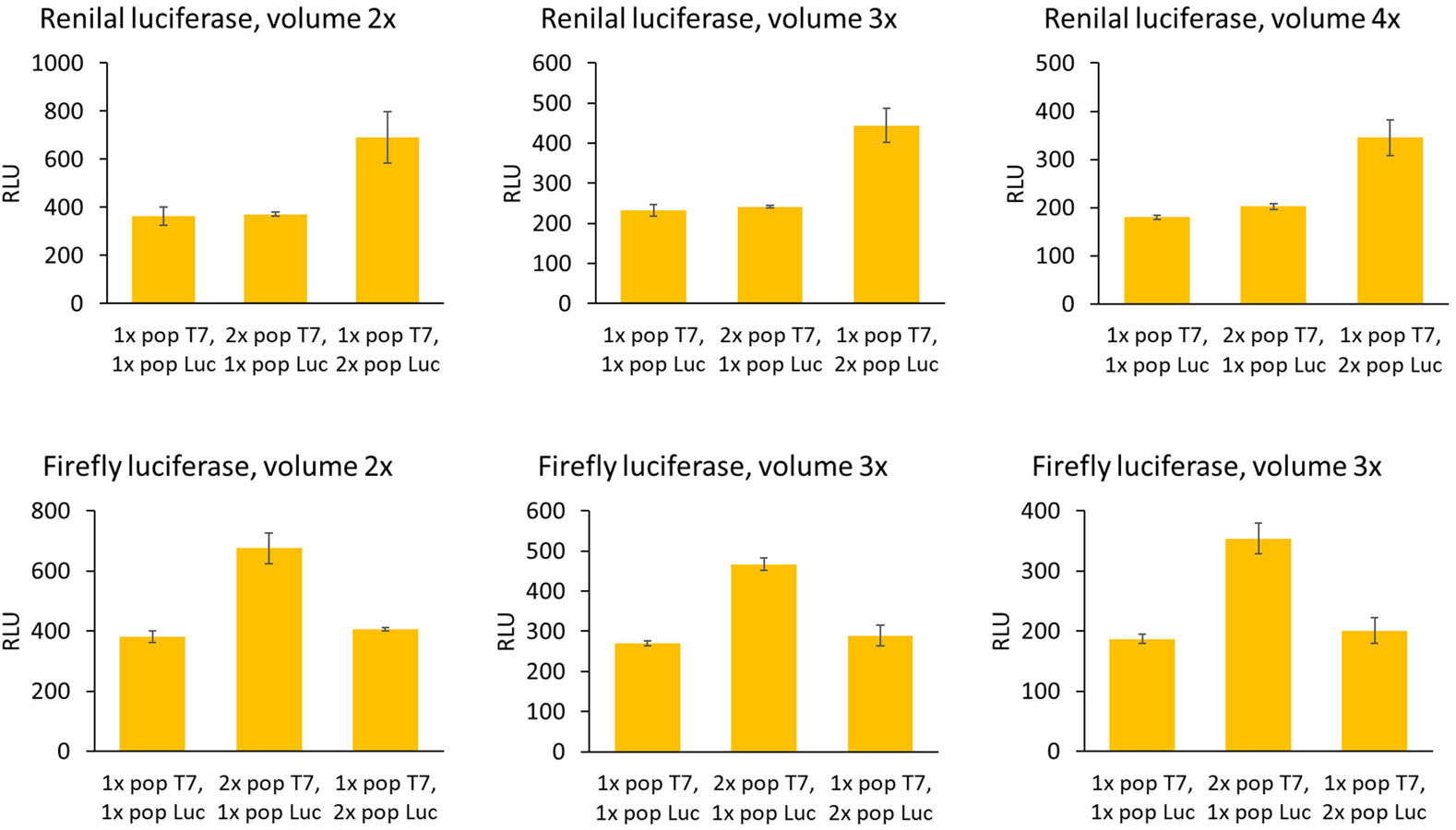
Individual experiment data for the heatmap showed on Figure 4e, Firefly luciferase and Renilla luciferase. Error bars indicate SEM, n=3.

**Figure S9.**
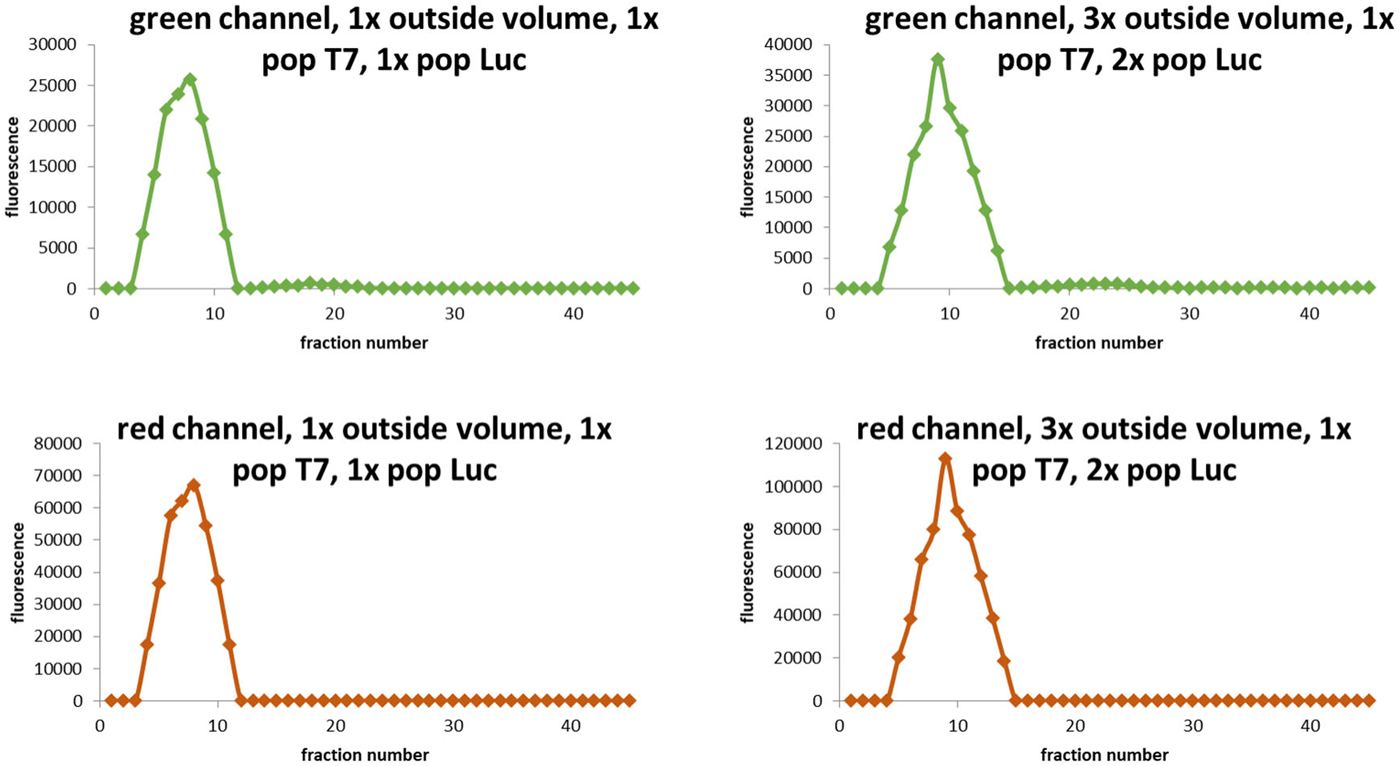
Size exclusion purification of samples with calcein inside synthetic cell liposomes (green channel), with membrane dyed with red Rhodamine dye (red channel). Those experiments confirm stability of synthetic cell liposomes of “Population Luc” after 16h incubation and the import of CPP tagged T7RNAP.

**Figure S10.**
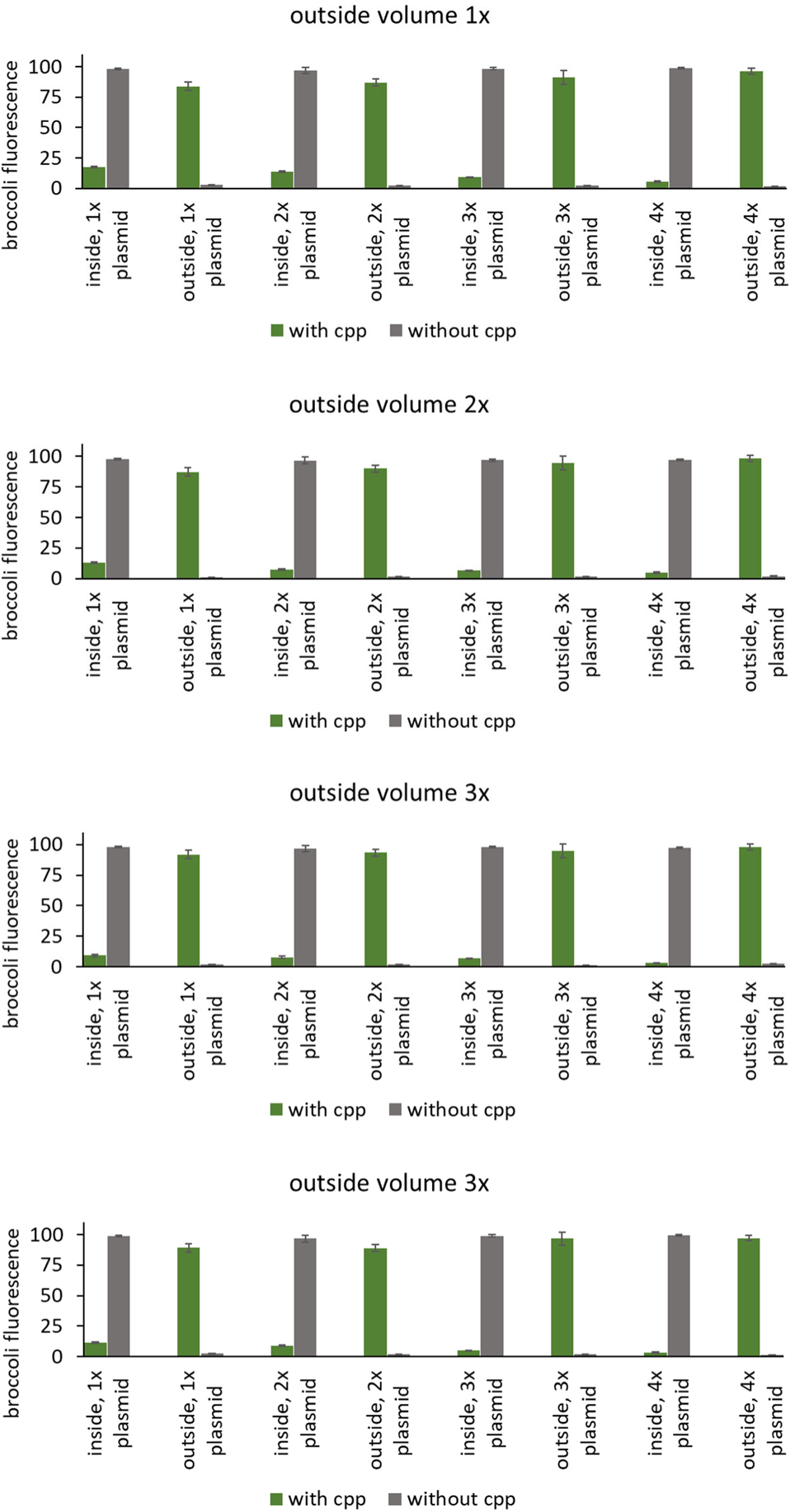
Individual experiments for the heatmap showed on Figure 5b and c. Error bars indicate SEM, n=3.

**Figure S11.**
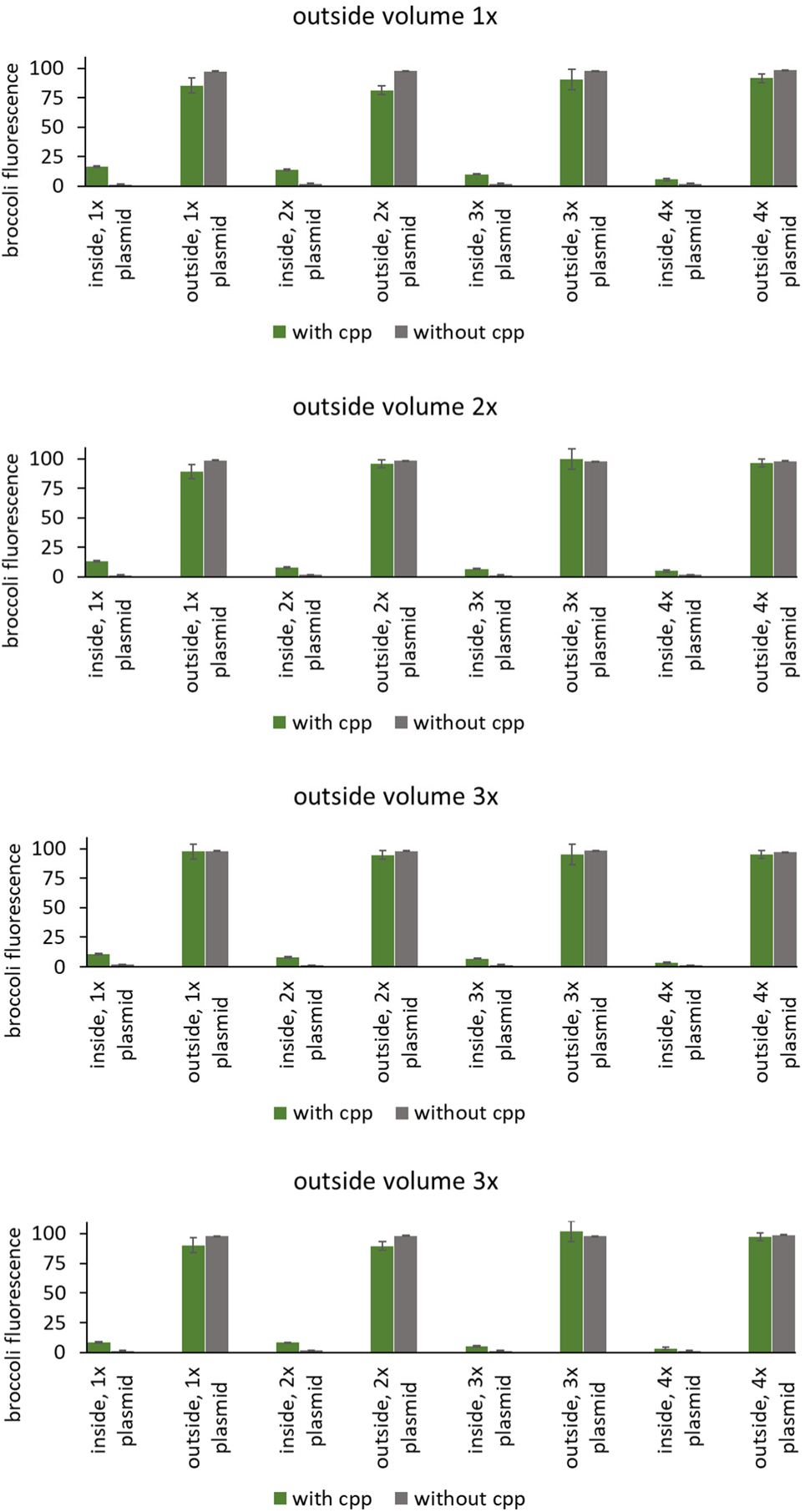
Individual experiments for the heatmap showed on Figure 5e and f. Error bars indicate SEM, n=3.

**Figure S12.**
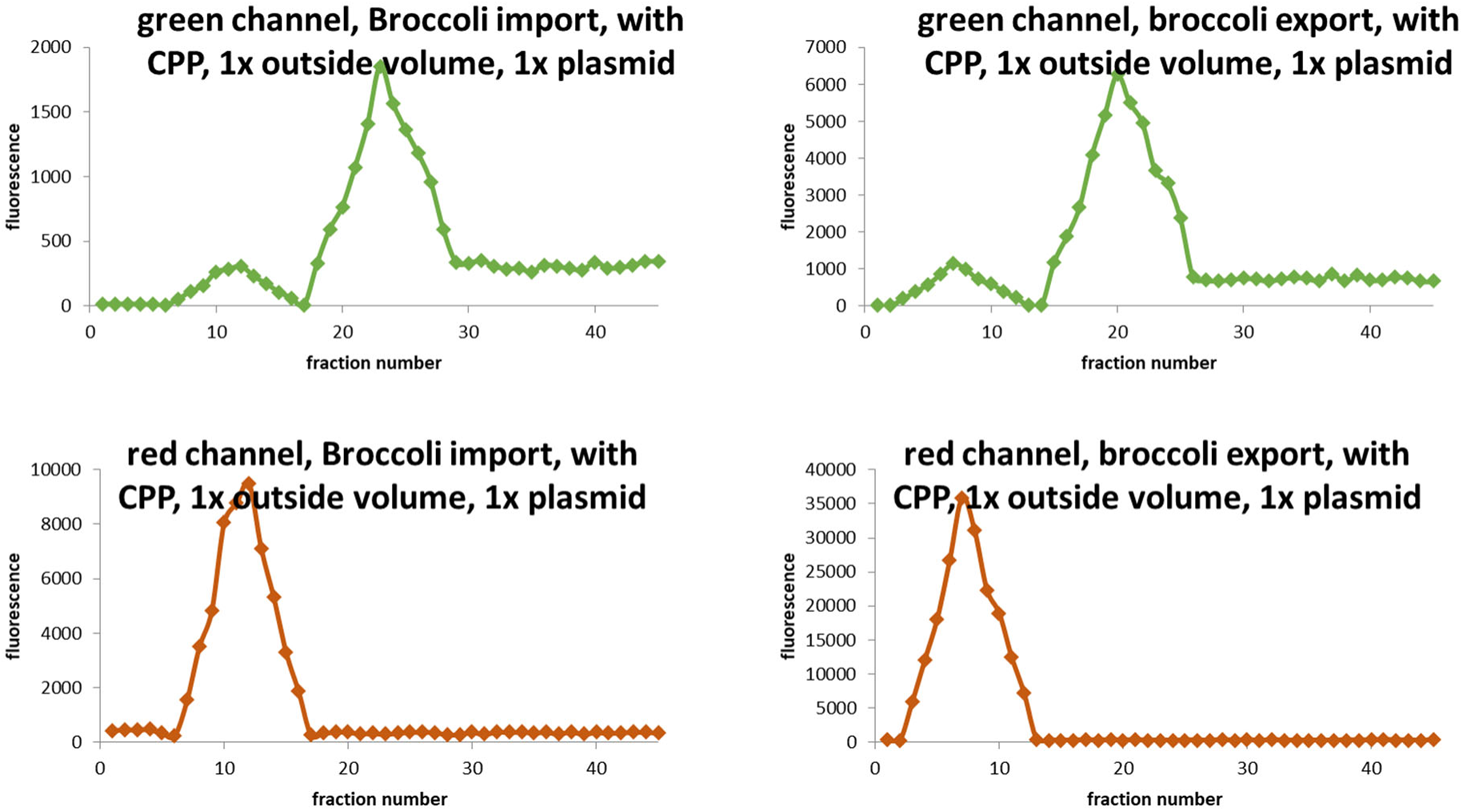
Size exclusion purification data. Synthetic cell liposomes were purified on Sepharose 4B. Fluorescence was measured in green channel (Broccoli) and red channel (Rhodamine membrane dye).

**Figure S13.**
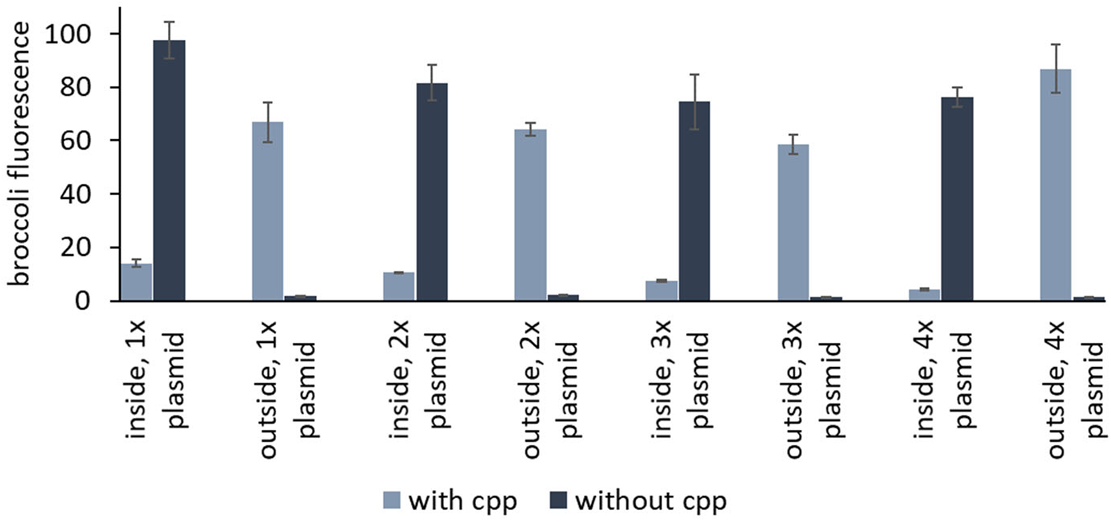
Export of RNA: Synthetic cells are filled with broccoli fluorescent RNA aptamer with MS2 RNA domain. CPP-tagged MS2 RNA binding domain is expressed outside of the cells. CPP-MS2 protein translocates inside cells, binds broccoli-MS2 RNA and translocates outside of the cell. The experimental workflow is the same as on Figure 5, except the translation of CPP-MS2 is done outside of the liposome. Error bars indicate S.E.M., n=3.

**Figure S14.**
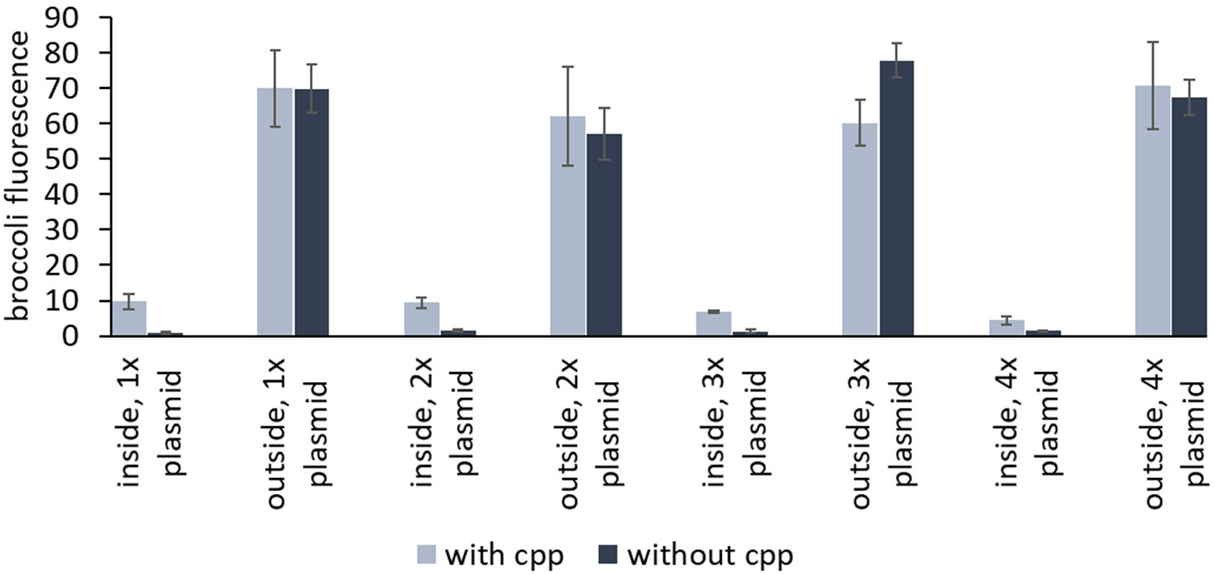
Import of RNA: Broccoli fluorescent RNA aptamer with MS2 RNA domain is added outside of liposomes (same liposomes as in synthetic cell experiments, but liposomes are filled with only Hepes buffer, no translation system, so they are not synthetic cells). CPP-tagged MS2 RNA binding domain is expressed outside of the liposome. CPP-MS2 protein binds broccoli-MS2 RNA and translocates inside liposome. The experimental workflow is the same as on Figure 5, except the translation of CPP-MS2 is done outside of the liposome. Error bars indicate S.E.M., n=3.

**Figure S15.**
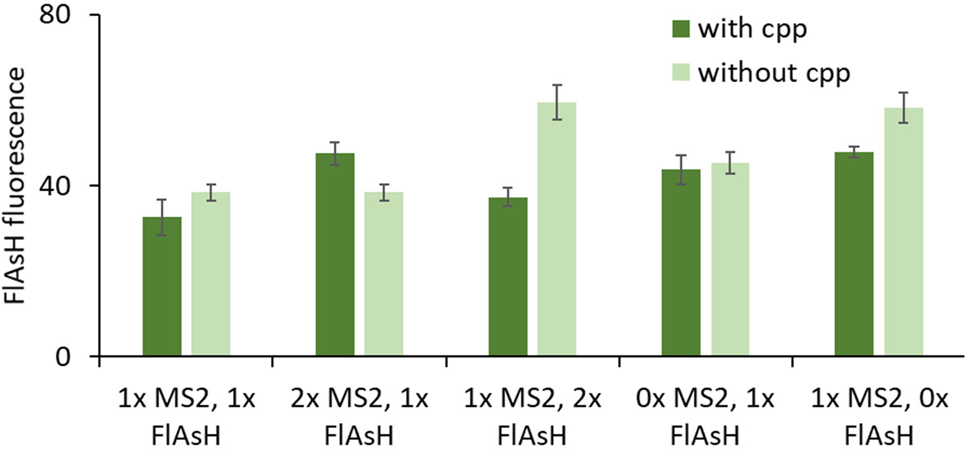
FlAsH fluorescence from samples without MS2 tag on FlAsH mRNA for experiment shown on figure 7, for different ratio of populations containing MS2 and FlAsH ligand. Fluorescence measured after 16h incubation. Error bars indicate S.E.M., n=3.

**Figure S16.**
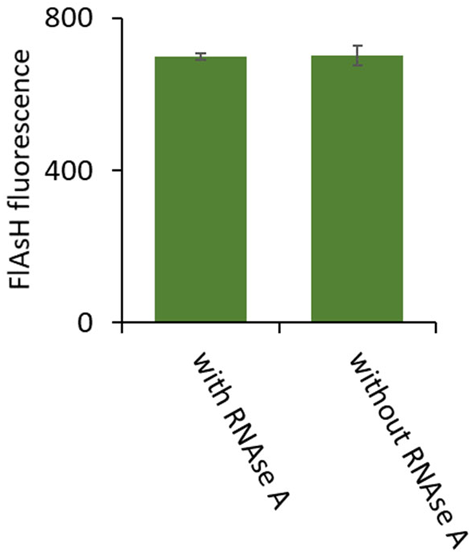
RNAse A is not permeable to synthetic cell membranes. Synthetic cells were prepared with FlAsH peptide mRNA and with FlAsH ligand, RNAse A was added to the outside of the cell. Fluorescence of FlAsH peptide was measured after 16h incubation. Error bars indicate S.E.M., n=3.

**Table S1.** Liposome concentrations corresponding to different lumen volume for 1uM POPC vesicles.

| Lipid concentration | Approximate lumen as %<br>volume of the whole<br>sample |
| --- | --- |
| 2mM | 7% |
| 1mM | 3.5% |
| 0.66mM | 2.3% |
| 0.5mM | 1.8% |

